# Vitamin B12 produced by gut bacteria modulates excitatory neurotransmission

**DOI:** 10.1101/2022.09.06.506833

**Authors:** Woo Kyu Kang, Antonia Araya, Bennett W. Fox, Andrea Thackeray, Frank C. Schroeder, Albertha J.M. Walhout, Mark J. Alkema

## Abstract

A growing body of evidence indicates that gut microbiota influence brain function and behavior. However, the molecular basis of how gut bacteria modulate host nervous system function is largely unknown. Here we show that vitamin B12-producing bacteria that colonize the intestine can modulate excitatory synaptic transmission and behavior in the host *Caenorhabditis elegans*. We find that vitamin B12 reduces cholinergic signaling in the nervous system through rewiring of the methionine (Met)/S-Adenosylmethionine (SAM) cycle in the intestine. We identify a conserved metabolic crosstalk between the Met/SAM cycle and the choline oxidation pathway. We show that metabolic rewiring of these pathways by vitamin B12 reduces cholinergic transmission by limiting the availability of free choline required by neurons to synthesize acetylcholine. Our study reveals a gut-brain communication pathway by which enteric bacteria modulate host behavior and may affect mental health.

## INTRODUCTION

There is growing interest in the relationship between gut microbiota, brain function and neurological disorders (Cryan et al., 2019). Changes in gut microbiota have been linked to several neurological conditions such as autism, anxiety, depression, schizophrenia, migraine, and neurodegeneration (Mohajeri et al., 2018; Sandhu et al., 2017; Zhu et al., 2020). However, it remains largely unclear whether imbalances in gut microbiota contribute to the neurological disorders or whether this dysbiosis is a consequence of the disorder. While probiotics and psychobiotics are being explored to improve physiology and neural health, the cause and effect relationships are hard to untangle (Mitrea et al., 2022). Often, the effect of diet and microbiota on health and behavior are subtle and only become apparent when the condition of the host is challenged (Buffington et al., 2021; Ezra-Nevo et al., 2020; Karl et al., 2018; Montgomery et al., 2020; Org et al., 2015). Elucidating the molecular mechanisms underlying the effects of gut microbiota on host nervous system function poses several additional challenges. The human microbiome consists of trillions of microorganisms and hundreds different bacterial species (Lloyd-Price et al., 2016) that could affect brain function of the host by modulating immune responses and through the production of highly diverse set of metabolites, and neurochemicals (Chow and Mazmanian, 2010; Fung et al., 2017). Even when potential beneficial microbiota are identified, the complexity of the mammalian nervous system and the gut microbiome makes it exceedingly difficult to determine the effects of specific bacterial species on neural function (Fischbach, 2018; Walter et al., 2020).

The nematode *C. elegans* provides a powerful system to study the microbiota-gut-brain interactions because its nervous system, and diet are relatively simple, well defined, and genetically tractable. In the wild, *C. elegans* feeds on diverse bacterial communities that can colonize its intestine (Berg et al., 2016; Dirksen et al., 2016; Samuel et al., 2016). In the laboratory *C. elegans* can be maintained on monoxenic bacterial culture, which simplifies mechanistic studies of microbial impacts on the host (Watson et al., 2014; Zhang et al., 2017). Several studies have begun to investigate the effects of specific bacteria and their metabolites on *C. elegans* physiology, lifespan, and behavior (Donato et al., 2017; Goya et al., 2020; O’Donnell et al., 2020; Seth et al., 2019; Smolentseva et al., 2017). However, the molecular mechanisms by which commensal gut bacteria modulate host neural function remain largely unknown.

In this study, we used *unc-2/CaV2*α*(gf)* mutants as a sensitized genetic background to facilitate the identification of bacteria that affect host neural function. *unc-2(gf)* mutants are hyperactive due to a gain-of-function (gf) mutation in the presynaptic voltage-gated calcium channel UNC-2/CaV2α (Huang et al., 2019). Similar mutations in the human CACNA1A/CaV2.1α channel cause familial hemiplegic migraine type 1 (FHM1) in human. The *unc-2(gf)* mutation, like CACNA1A FHM1 mutations, causes increased excitatory transmission and excitatory/inhibitory imbalance (Huang et al., 2019). Other neurological disorders with excitation-inhibition imbalance, such as autism, epilepsy, and migraine, have been associated with alterations in the gut microbiota (Arzani et al., 2020; Dahlin and Prast-Nielsen, 2019; Golubeva et al., 2017; Lum et al., 2020; Sharon et al., 2019; van Hemert et al., 2014). We find that vitamin B12-producing bacteria that colonize the intestine modulate excitatory synaptic transmission of *C. elegans*. We show that vitamin B12 reduces cholinergic transmission in the nervous system through metabolic crosstalk between vitamin B12-dependent Met/SAM cycle and the choline oxidation pathway in the intestine. We find that vitamin B12 drives metabolic rewiring and reduces the availability of free choline which is taken up by neurons for the synthesis of acetylcholine. Our study reveals a molecular mechanism of gut-brain communication by which gut microbiota modulate host nervous system function and behavior and may explain the positive impact of vitamin B12 on neural disorders.

## RESULTS

### Vitamin B12 producing bacteria suppress hyperactive behavior of *unc-2(gf)* mutants

We surveyed bacterial diets for their ability to suppress the hyperactive behavior of *unc-2(gf)* mutants. *unc-2(gf)* mutant animals have an increased locomotion rate and display a striking clonic seizure-like behavior characterized by frequent jerky reversals (Huang et al., 2019). This behavioral hyperactivity can be easily quantified by the scoring reversal frequency with an automated multi-worm tracking system (Swierczek et al., 2011) and provides a simple and sensitive readout to screen the impact of bacterial diets on host neural function. To rule out potential confounding effects of the bacterial diet on neuronal development, animals were grown on a standard laboratory *E. coli* OP50 diet until the late 4th larval stage (L4). L4 were then transferred to plates with single bacterial diets. After 24h, young adult *unc-2(gf)* animals were transferred to a thin lawn of *E. coli* OP50, to quantify reversal frequency. We tested 40 bacterial strains of 20 different genera, most of which have been found to be associated with *C. elegans* in the wild (Berg et al., 2016; Dirksen et al., 2016; Samuel et al., 2016). The selected strains supported normal animal growth and development and are non-pathogenic (Figure S1A and S1B).

We found that 18 of the 40 bacterial diets significantly suppressed the hyper-reversal behavior of *unc-2(gf)* mutants compared to the standard *E. coli* OP50 diet (Figure 1A). Unlike *E. coli*, some of the bacterial species that suppressed hyper-reversal behavior, such as *Comamonas aquatica* and *Pseudomonas putida*, are known to produce vitamin B12 (Martens et al., 2002; Watson et al., 2014). Since vitamin B12 plays a crucial role in health and neural function (Green et al., 2017), we analyzed all bacteria for vitamin B12 production using a *C. elegans Pacdh-1::GFP* reporter strain. *acdh-1::GFP* expression is strongly repressed by the presence of dietary vitamin B12 (Arda et al., 2010; Watson et al., 2014). All bacterial strains that repressed *Pacdh-1::GFP* expression also suppressed *unc-2(gf)* reversals (Figures 1A and 1B; Figure S1C). This suggests that vitamin B12 could be a key bacterial metabolite that suppresses the hyper-reversal behavior of *unc-2(gf)* mutants. This is further supported by the observation that a *C. aquatica cbiAΔ* strain, that cannot produce vitamin B12 (Watson et al., 2014), failed to reduce reversals of *unc-2(gf)* mutants (Figure 1C and Figure S1D).

**Figure 1.**
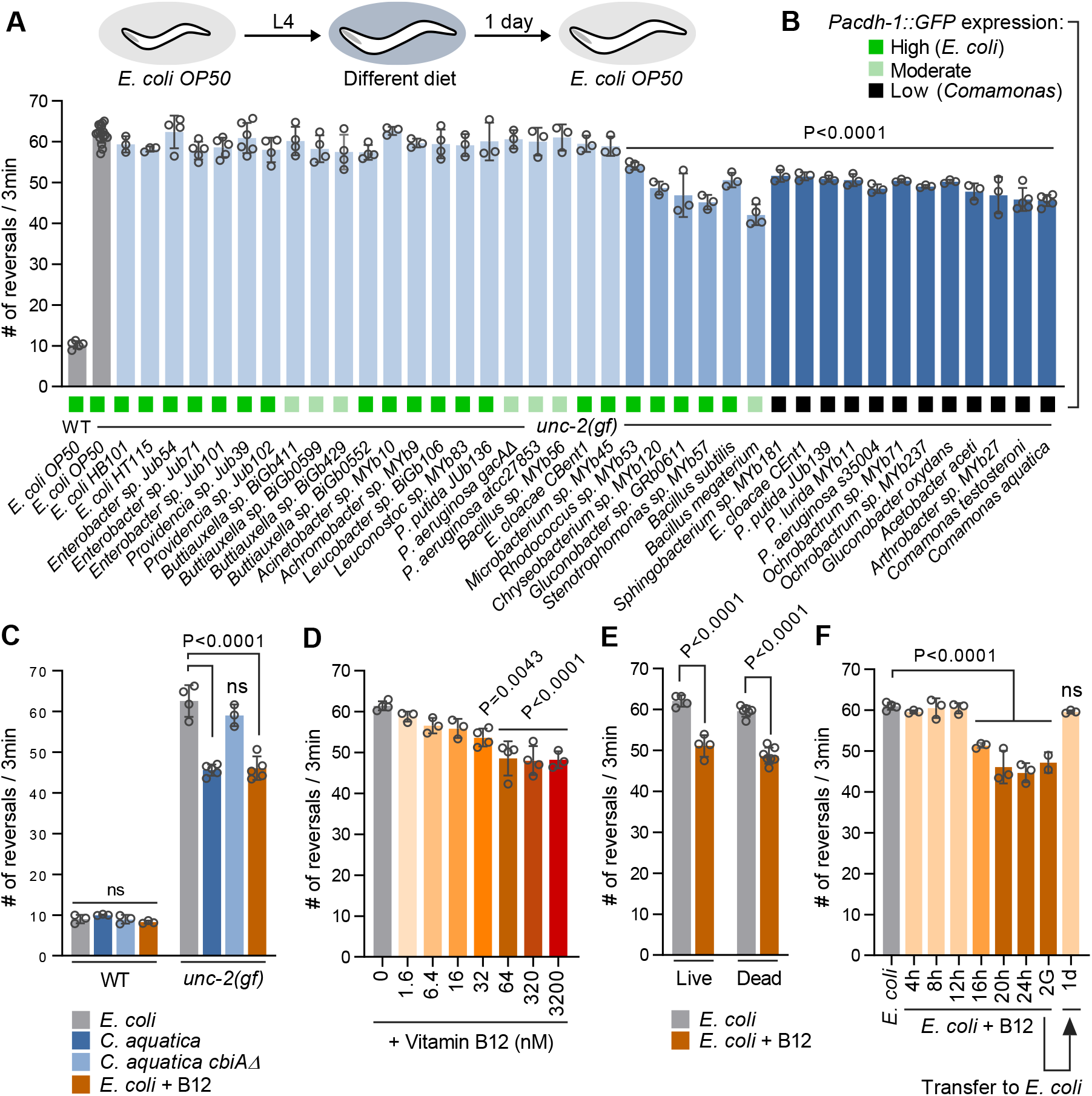
Vitamin B12 produced by gut bacteria suppresses hyperactive behavior of *unc-2(gf)* mutants. (A) Reversal frequency of *unc-2(gf)* mutants fed the indicated bacterial strains for 24 h. (B) GFP expression of *Pacdh-1::GFP* animals fed the indicated bacterial strains for 24 h. Shades of green represent relative GFP expression levels. (C) Reversal frequency of wild-type and *unc-2(gf)* mutants fed *E. coli, C. aquatica, C. aquatica cbiAΔ* (B12-), or *E. coli* supplemented with 64 nM B12. (D) Reversal frequency of *unc-2(gf)* mutants fed *E. coli* supplemented with the indicated concentrations (nM) of B12. (E) Reversal frequency of *unc-2(gf)* mutants fed live and heat-killed *E. coli* with or without 64 nM B12. (F) Reversal frequency of *unc-2(gf)* mutants fed *E. coli* supplemented with 64 nM B12 for the indicated time and after transfer to *E. coli*. All data are presented as mean ± s.e.m. from at least three independent experiments. P-values are calculated by one-way ANOVA with Dunnett’s multiple comparison test; ns, not significant. See also Figure S1.

### Gut colonization by *Comamonas* modulates *C. elegans* behavior

To determine whether *Comamonas* establishes a gut microbiome, we examined the ability of *C. aquatica* to colonize the intestine of *C. elegans*. L4 animals were transferred from *E*.*coli* OP50 diet to different bacterial diets for 24h after which the *C. elegans* were homogenized and the number of colony forming units (CFUs) per worm calculated (Berg et al., 2016; Dirksen et al., 2016). *C. aquatica* efficiently colonized the intestine of worms with a density of ∼ 2 × 10^3^ CFUs per animal, compared to ∼ 80 CFUs for *E. coli* OP50 and ∼ 2.7 × 10^4^ CFUs for the pathogenic bacteria *Pseudomonas aeruginosa* (Figure 2A). *C. aquatica* colonization was stable for at least 3 days after a transfer to an *E. coli* diet (Figure 2B). Consistently, we found that *unc-2(gf)* mutant animals grown on live *C. aquatica* still showed reduced reversals three days after a shift to an *E. coli* diet (Figure 2C). In contrast, sustained reversal suppression was not observed when *unc-2(gf)* mutants were fed killed *C. aquatica* (Figure 2C). Thus, persistent changes in the behavior correlate with the presence of *C. aquatica* in the *C. elegans* intestine. To further test whether gut colonization by *C. aquatica* can drive reversal suppression, we treated worms raised on live *C. aquatica* with antibiotics. Kanamycin treatment (200 µg/ml) effectively eliminated *C. aquatica* from the *C. elegans* gut (Figure 2D; Figures S2A–S2B). The reversal suppression of *unc-2(gf)* mutants colonized with *C. aquatica* was no longer observed after clearance of gut bacteria with kanamycin treatment (Figure 2E). The effect of Kanamycin was specific to *C. aquatica* as the treatment did not affect the reversal behavior of *unc-2(gf)* mutants fed *E. coli* OP50 (Figures S2C). Together, these results indicate that *C. aquatica* colonization of the intestine and lead to persistent changes in behavior.

**Figure 2.**
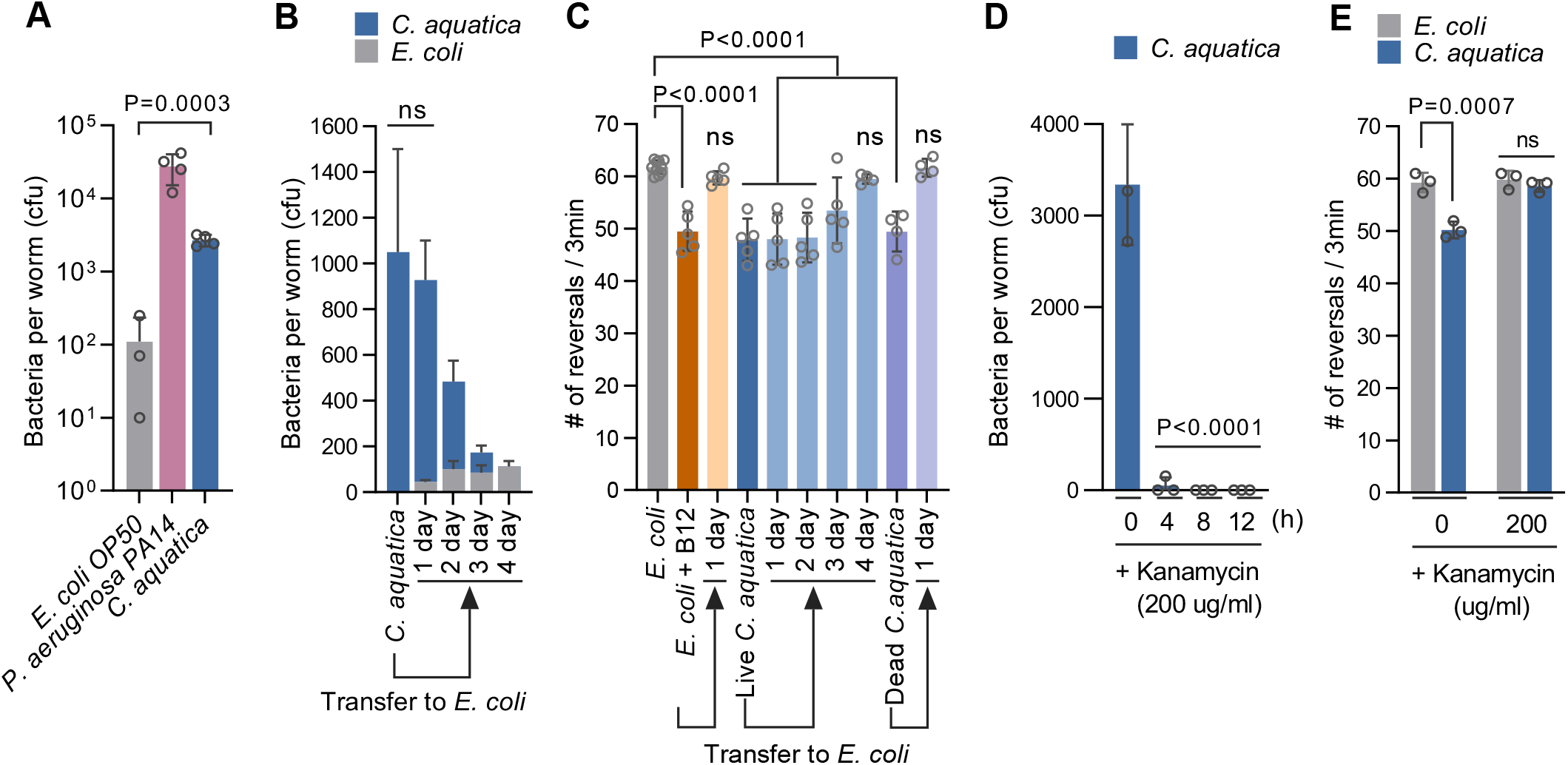
*Comamonas* colonizes the *C. elegans* intestine and modulates behavior. (A) Bacterial colony forming units (CFU) per animal from *unc-2(gf)* mutants fed *E. coli, P. aeruginosa PA14*, or *C. aquatica*. (B) Bacterial CFU per animal from *unc-2(gf)* mutants fed *C. aquatica* after transfer to *E. coli*, measured over multiple days, as indicated. (C) Reversal frequency of *unc-2(gf)* mutants fed *E. coli* with 64 nM B12, *C. aquatica*, or dead *C. aquatica* after transfer to *E. coli*, measured over multiple days, as indicated. (D) Bacterial CFU per animal from *unc-2(gf)* mutants fed *C. aquatica* treated with 200 µg/ml kanamycin for the indicated time. (E) Reversal frequency of *unc-2(gf)* mutants colonized with *C. aquatica* after kanamycin treatment for 24 h. All data are presented as mean ± s.e.m. from at least three independent experiments. P-values are from one-way ANOVA with Dunnett’s multiple comparison test; ns, not significant. See also Figure S2.

### Vitamin B12 supplementation is sufficient to suppress hyperactive behavior of *unc-2(gf)* mutants

Vitamin B12 is exclusively synthesized by bacteria and archaea (Bender, 2003). Animals acquire B12 through symbiotic relationship with bacteria in their gut or their diet (LeBlanc et al., 2013; Watanabe and Bito, 2018). *E. coli* can import vitamin B12 from regular growth media but the concentrations are much lower than those found in B12 producing *Comamonas* (Bito et al., 2013; Watson et al., 2014). We tested whether dietary vitamin B12 is sufficient to change behavior. Supplementation of vitamin B12 to *E. coli* (> 64 nM) suppressed the reversals of *unc-2(gf)* mutants to a similar extent as *C. aquatica* (Figures 1C and 1D; Figure S1D). Vitamin B12 reduced the reversals of *unc-2(gf)* mutants equally well, when supplemented to either live or dead *E. coli* (Figure 1E and Figure S1E). This indicates that the B12 effect was not mediated by secondary alterations in bacterial metabolism and is sufficient to change host behavior. The suppression effect of dietary B12 supplementation on reversal behavior took effect within 16 h (Figure 1F). Reversal suppression upon B12 supplementation disappeared a day after transfer to a regular *E. coli* diet without vitamin B12 (Figure 1F). Notably, the reversal suppression of an *E. coli* diet supplemented with B12 persisted markedly shorter than that of live *C. aquatica* diet (Figure 2C). This further supports our observation that *Comamonas* gut colonization leads to persistent behavioral changes of the host.

### Vitamin B12 inhibits excitatory cholinergic neurotransmission

How does vitamin B12 impact neural function and behavior? The *unc-2(gf)* mutants used in our screen have increased excitatory cholinergic neurotransmission (Huang et al., 2019). Consequently, *unc-2(gf)* mutants are hypersensitive to the acetylcholine esterase inhibitor, aldicarb. In the presence of aldicarb, acetylcholine accumulates at the neuromuscular junction, which leads to muscle hyper-contraction and eventual paralysis (Nguyen et al., 1995). Wild-type animals completely paralyzed on in 1 mM aldicarb plates within 100 min. In contrast, *unc-2(gf)* mutants that were raised on an *E. coli* diet completely paralyzed in 1 mM aldicarb within 40 min. B12-supplemented *E. coli* diet significantly slowed (60 min) aldicarb induced paralysis of *unc-2(gf)* animals (Figure 3A), suggesting that vitamin B12 inhibits cholinergic transmission in *unc-2(gf)* mutants. To exclude the possibility that B12 affects aldicarb uptake, we analyzed whether B12 affects the severe locomotion defects of animals lacking the acetylcholinesterase *ace-1* and *ace-2* genes (Culotti et al., 1981). Consistent with our aldicarb data, we found that a B12 supplemented *E. coli* diet or a *C. aquatica* diet dramatically improved the severe locomotion defect of *ace-1; ace-2* double mutants (Culotti et al., 1981) (Figures 3B and 3C; Figure S3A).

**Figure 3.**
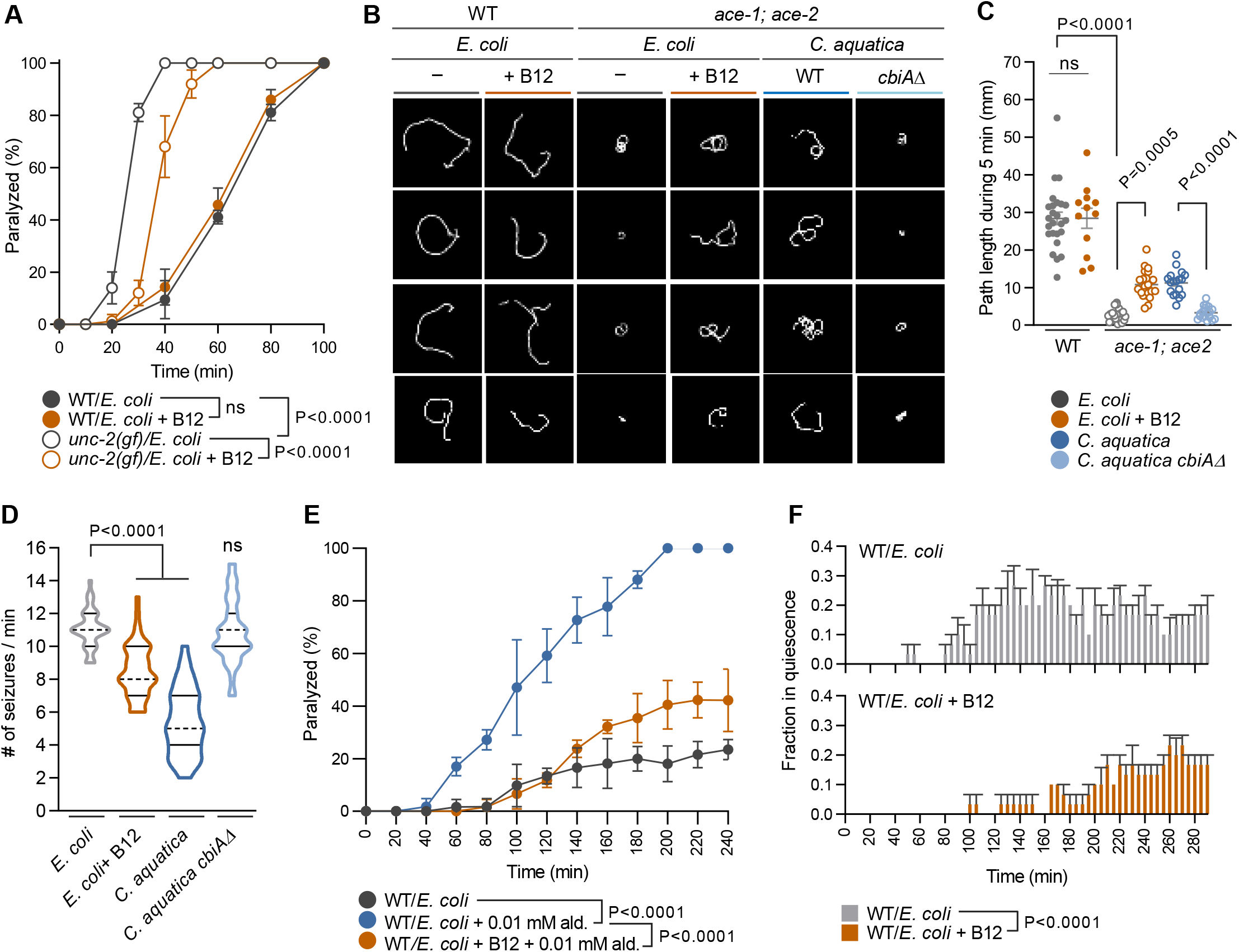
Vitamin B12 inhibits cholinergic transmission. (A) Percentage of paralyzed wild-type and *unc-2(gf)* mutants fed *E. coli* with or without 64 nM B12 on 1 mM aldicarb. (B and C) Tracks (B) and quantification of path length (C) of single young adult animal of wild-type and *ace-1; ace-2* double mutants fed *E. coli, E. coli* supplemented with 64 nM B12, *C. aquatica*, or *C. aquatica cbiAΔ* for 10 min. (D) Quantification of convulsion phenotype of *acr-2(gf)* mutants fed *E. coli, E. coli* supplemented with 64 nM B12, *C. aquatica*, or *C. aquatica cbiAΔ* for 24 h. (E) Percentage of paralyzed wild-type animals fed *E. coli* with or without 64 nM B12 on 0.01 mM aldicarb during swimming. (F) Vitamin B12 reduces swimming-induced quiescence. Fraction of wild-type animals fed *E. coli* with or without 64 nM B12 in quiescence. All data are presented as mean ± s.e.m. from at least three independent experiments. P-values are from two-way ANOVA with Tukey’s multiple comparison test; ns, not significant. See also Figure S3.

To further study the effect of vitamin B12 on excitatory synaptic transmission, we analyzed additional mutants with increased cholinergic signaling. Mutants with a gain-of-function mutation in the ACR-2 acetylcholine receptor subunit *(acr-2(gf))* mutants display epileptic-like convulsive muscle contractions, as the result of increased endogenous cholinergic excitation and reduced GABAergic inhibition in the motor circuit. The imbalance of excitation and inhibition (E/I) in *acr*-*2(gf)* mutants resembles that reported for gain-of-function mutations of nicotinic acetylcholine receptors (nAChRs) in frontal lobe epilepsy (Becchetti et al., 2015). We found that a diet of either B12 supplemented *E. coli* or a *C. aquatica* significantly reduced convulsive muscle contractions of the *acr-2(gf)* mutant animals, whereas a diet of mutant *C. aquatica cbiAΔ* failed to do so (Figure 3D). Together, these data indicate that vitamin B12 reduces cholinergic signaling in mutants with E/I imbalance.

### Vitamin B12 reduces swimming induced quiescence

In contrast to the cholinergic signaling mutants, vitamin B12 had no obvious effects on locomotion behavior or aldicarb sensitivity of wild-type animals under standard conditions (Figure 3A; Figures S3B–S3E). Since the effects of gut microbiota and individual nutrients can be revealed by environmental and genetic conditions of the host, we hypothesized that the effect of vitamin B12 in wild-type animals may become more apparent under conditions of increased cholinergic signaling. Cholinergic transmission controls the body wall muscle contractions that propel worm movement. When *C. elegans* transitions from a solid agar surface to liquid, it dramatically increases its undulation frequency as it switches from crawling to swimming (Pierce-Shimomura et al., 2008). When exposed to 1 mM aldicarb in liquid, swimming animals became paralyzed within 20 min; compared to 100 min when crawling on agar plates (Figure S3F). The enhanced aldicarb sensitivity of wild-type animals in liquid is consistent with increased cholinergic signaling during swimming. We observed that vitamin B12 supplementation significantly suppressed the aldicarb sensitivity of swimming wild-type animals (Figure 3E). High levels of cholinergic signaling has also been shown to induce bouts of quiescent behavior characterized by transient paralysis during prolonged swimming in liquid (Ghosh and Emmons, 2008). We found that supplementation of vitamin B12 also strongly suppressed the swimming-induced quiescence in wild-type animals (Figure 3F). Together, these results suggest that vitamin B12 reduces cholinergic transmission in wild-type animals under conditions that increase acetylcholine release.

### Vitamin B12 modulates cholinergic signaling through its role in the Met/SAM cycle in the intestine and hypodermis

Vitamin B12 is an essential cofactor for two metabolic enzymes in both *C. elegans* and mammals (Watson et al., 2014): methionine synthase which converts homocysteine to methionine in the Methionine/S-Adenosylmethionine (Met/SAM) cycle (*metr-1*; Figure 4A) and methylmalonyl-CoA mutase which is required for propionyl-CoA breakdown (*mmcm-1*; Figure 4B). We generated *metr-1; unc-2(gf) and mmcm-1; unc-2(gf)* double mutants to determine whether either of these B12 dependent pathways modulates cholinergic signaling. The suppression effect of vitamin B12 on the reversals of *unc-2(gf)* animals was completely abolished by a *metr-1* mutation, but not by an *mmcm-1* mutation (Figure 4C). Notably, the *metr-1* mutation further increased reversals of *unc-2(gf)* mutants, whereas the *mmcm-1* mutation had no such effect (Figure 4C). Vitamin B12 also failed to suppress reversal frequency by mutations in *sams-1*, the predicted *C. elegans* SAM synthetase in the Met/SAM cycle, but not by mutations in *pcca-1* or *mce-1*, which are related to the propionyl-CoA metabolic pathway (Figures S4A and S4B). Furthermore, the effect of vitamin B12 on aldicarb sensitivity was completely abolished in *metr-1; unc-2(gf)* (Figure 4D) and *sams-1; unc-2(gf)* mutants (Figure S4C). *metr-1* single mutants also displayed a marked increase in swimming-induced quiescence (Figure 4E). In contrast wild-type animals, vitamin B12 supplementation failed to suppress quiescence in the *metr-1* mutants (Figure 4E). This indicates that the effect of vitamin B12 on behavior depends on *metr-1* and the Met/SAM cycle.

**Figure 4.**
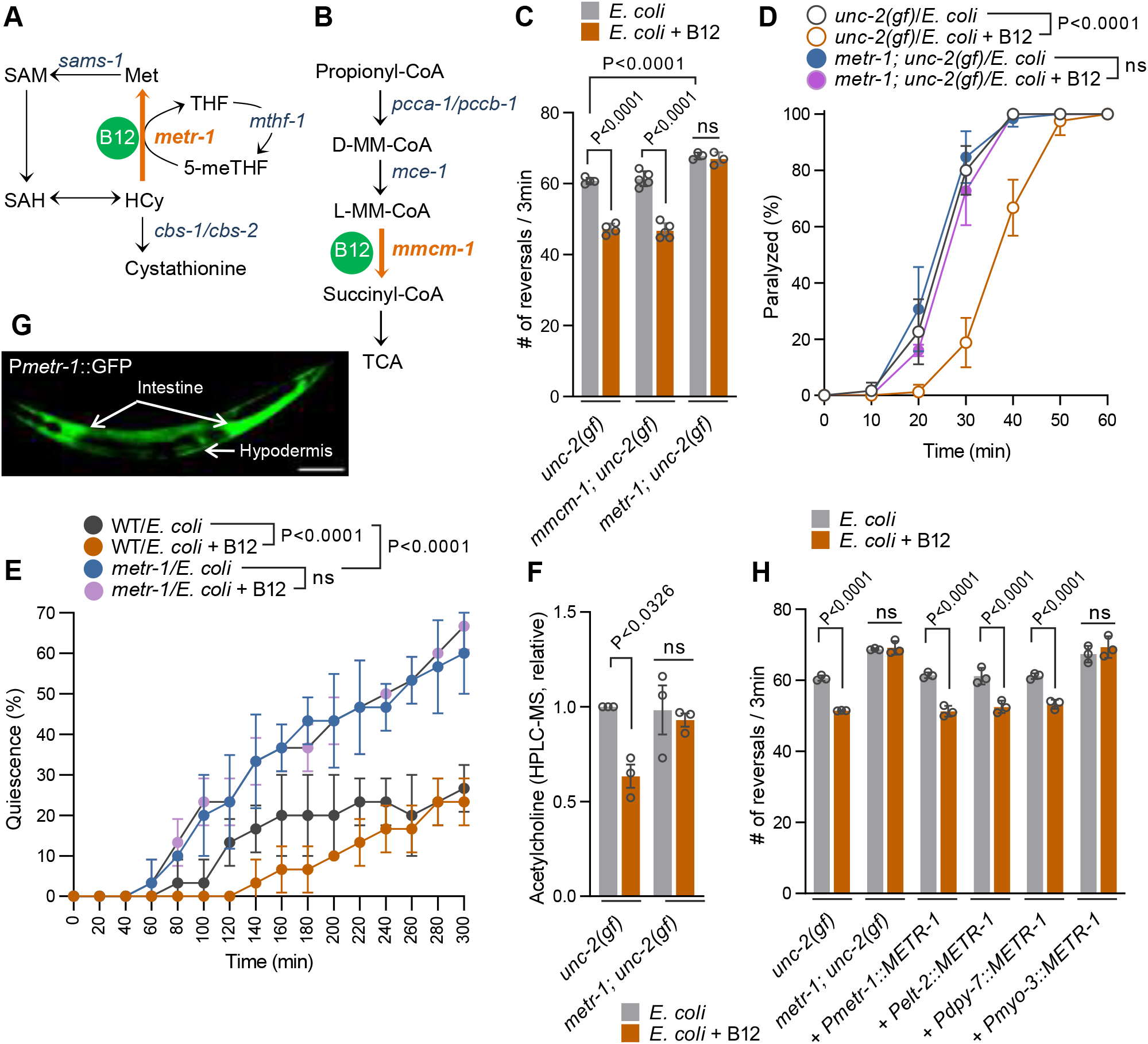
Vitamin B12 dependent Met/SAM cycle acts in intestine and hypodermis to modulate behavior. (A and B) B12-dependent metabolic pathways, Met/SAM cycle (A) and propionyl-CoA breakdown pathway (B) are highly conserved in *C. elegans*. (C) Reversal frequency of *metr-1; unc-2(gf)* or *mmcm-1; unc-2(gf)* mutants fed *E. coli* with or without 64 nM B12. (D) Percentage of paralyzed *metr-1; unc-2(gf)* mutants fed *E. coli* with or without 64 nM B12 on 1 mM aldicarb. (E) Fraction of wild-type and *metr-1* mutants fed *E. coli* with or without 64 nM B12 in quiescence. (F) Quantification of acetylcholine in *unc-2(gf)* and *metr-1; unc-2(gf)* mutants fed *E. coli* with or without 64 nM B12 for 24 h. (G) Expression pattern of *Pmetr-1::GFP*. Scale bar is 100 μm. (H) Reversal frequency of *metr-1; unc-2(gf)* mutant animals expressing *metr-1* cDNA driven by *Pmetr-1* (endogenous), *Pelt-2* (intestinal), *Pdpy-7* (hypodermal), or *Pmyo-3* (muscle) promoter fed *E. coli* with or without 64 nM B12. All data are presented as mean ± s.e.m. from at least three independent experiments. P-values are from one-way ANOVA with Tukey’s multiple comparison test; ns, not significant. See also Figure S4.

Since vitamin B12 affected cholinergic transmission, we analyzed the effect of B12 supplementation on acetylcholine levels in whole worm extracts. We found that *unc-2(gf)* mutants had a marked reduction in acetylcholine levels in both the HPLC-MS analyses (37%) and fluorometric assays (28%) when *E. coli* diet was supplemented with B12 (Figure 4F and Figure S4D). In contrast, vitamin B12 supplementation did not affect acetylcholine levels in *metr-1; unc-2(gf)* mutants (Figure 4F). Taken together, these data indicate that vitamin B12 reduces cholinergic transmission through its role in the Met/SAM cycle.

To determine the site of action for the Met/SAM cycle, we analyzed the *metr-1* expression pattern using a *Pmetr-1::GFP* transcriptional reporter. The *Pmetr-1::GFP* fluorescence was observed in the intestine and hypodermis, but not in neurons (Figure 4G). We expressed wild-type *metr-1* cDNA under tissue-specific promoters in the *metr-1; unc-2(gf)* mutants. Expression of *metr-1* in the intestine or hypodermis, but not in muscle, was sufficient to restore the B12 dependent suppression of reversals in *metr-1; unc-2(gf)* mutants (Figure 4H and Figure S4E). This indicates that vitamin B12 modulates cholinergic transmission cell non-autonomously through its role as a co-factor METR-1 and the Met/SAM cycle in intestine and hypodermis.

Next, we asked whether B12 suppressed reversals of *unc-2(gf)* mutants as the result of accumulation or depletion of metabolites from the Met/SAM cycle. METR-1 catalyzes the conversion of homocysteine to methionine, which can be further converted to SAM by the S-Adenosylmethionine synthase SAMS-1 (Li et al., 2011; Walker et al., 2011) (Figure 4A). Supplementation of methionine or SAM increased animal growth consistent with previously reports (Watson et al., 2014), but did not suppress the reversal frequency of *unc-2(gf)* mutants (Figures S4F and S4G). Increased levels of homocysteine are associated with the symptoms of a variety of neurological disorders, including depression autism and migraine (Moustafa et al., 2014; Obeid et al., 2007; Puig-Alcaraz et al., 2015). This finding has led to the “homocysteine hypothesis” which proposes that B12 deficiency and increased homocysteine levels underlie the etiology of these disorders (Folstein et al., 2007). However, homocysteine supplementation did not block the effect of vitamin B12 on animal growth and reversal frequency of *unc-2(gf)* mutants (Figures S4F and S4G). Together, these results suggest that the effect of vitamin B12 on behavior is independent of its role in animal growth and may be mediated indirectly by increased metabolic flux through the Met/SAM cycle.

### Vitamin B12 modulates cholinergic signaling through metabolic crosstalk between the Met/SAM cycle and the choline oxidation pathway

How does the vitamin B12-dependent Met/SAM cycle in the intestine and hypodermis modulate cholinergic transmission in the nervous systems? In mammals, the Met/SAM cycle is tightly associated with the choline oxidation pathway (Mato et al., 2002; Shinohara et al., 2006; Xue and Snoswell, 1985). The choline oxidation pathway converts choline into betaine, which in turn can serve as a methyl donor for the synthesis of methionine in the Met/SAM cycle (Figure 5A) (da Costa et al., 2005; Ueland, 2011). Choline is also the rate-limiting precursor of acetylcholine biosynthesis in the nervous system (Birks and Fitch, 1974; Birks and Macintosh, 1957). To examine metabolic crosstalk between the Met/SAM cycle and cholinergic signaling, we tested the effect of exogenous choline on reversal behavior of *unc-2(gf)* mutants. Dietary choline supplementation completely abolished the effect of B12 on reversal suppression (Figure 5B) but had no effect on reversals in the absence of vitamin B12 (Figure 5B). Furthermore, vitamin B12 failed to suppress the swimming-induced quiescence in the presence of exogenous choline (Figure 5C). Choline supplementation did not affect vitamin B12’s positive impact on animal growth (Figure S5A), consistent with our finding that vitamin B12 suppresses reversal behavior independently of its role in animal growth. To determine if vitamin B12 affects choline levels, we measured the amount of free choline (non-bound forms) using an enzyme-based fluorometric assay. Vitamin B12 supplementation led to a 25% decrease of free choline levels in the *unc-2(gf)* mutants (Figure 5D).

**Figure 5.**
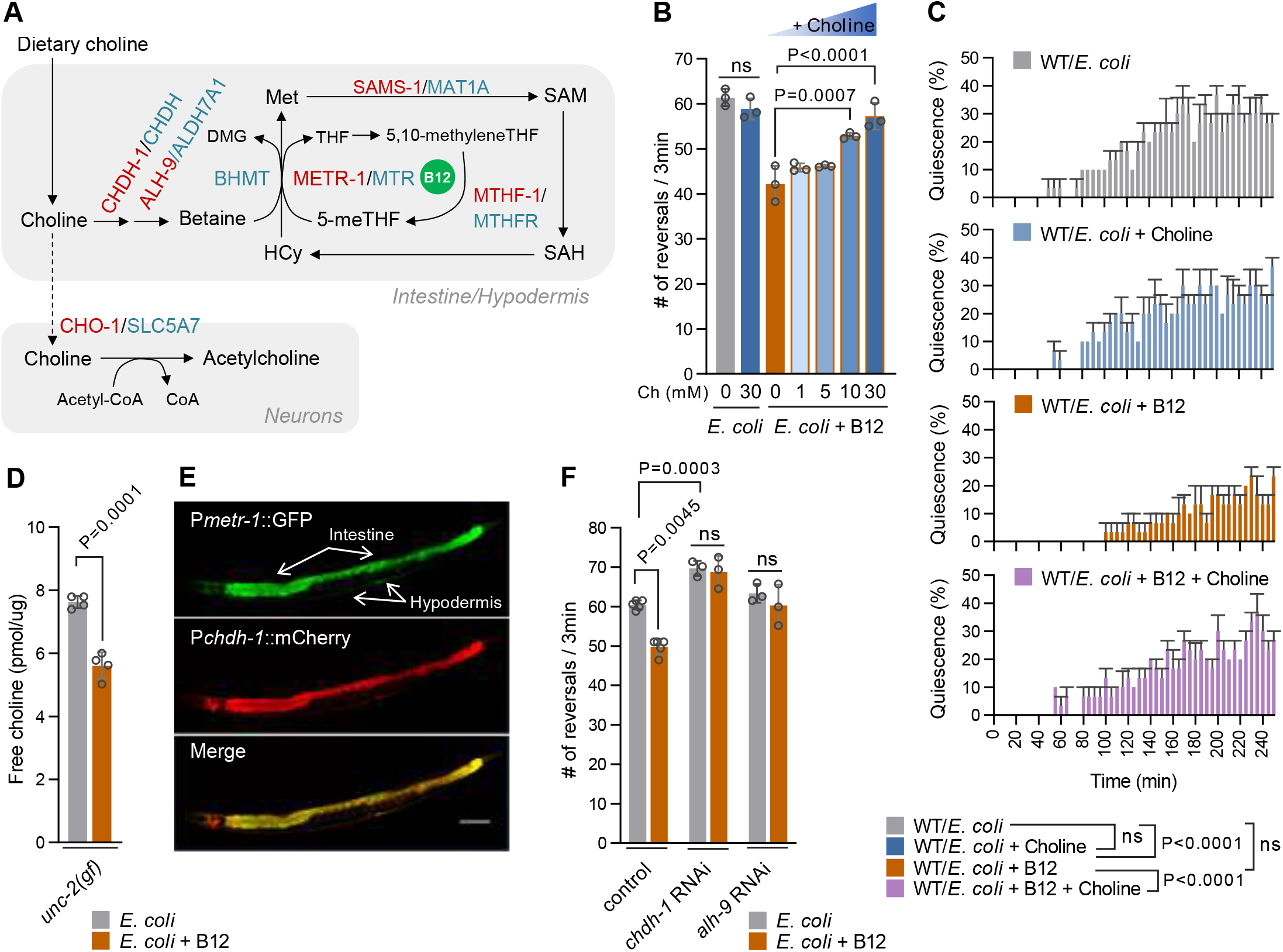
Crosstalk between Met/SAM cycle and choline metabolism modulates excitatory cholinergic transmission. (A) Metabolic network of Met/SAM cycle and choline metabolism. Red and blue indicate *C. elegans* and human metabolic enzymes involved, respectively. (B) Reversal frequency of *unc-2(gf)* mutants fed *E. coli* with or without 64 nM B12 with indicated concentrations of choline. (C) Fraction of wild-type animals fed *E. coli, E. coli* with choline (30 mM), *E. coli* with B12 (64 nM), or *E. coli* with both B12 and choline in quiescence. (D) Quantification of free choline of *unc-2(gf)* mutants fed *E. coli* with or without 64 nM B12 for 24 h. (E) Co-expression pattern of *Pmetr-1::GFP* and *Pchdh-1::mCherry* in the intestine and hypodermis. Scale bar is 100 µm. (F) Reversal frequency of *unc-2(gf)* mutants subjected to RNAi knockdown of *chdh-1* or *alh-9* fed *E. coli* with or without 64 nM B12. All data are presented as mean ± s.e.m. from at least three independent experiments. P-values are from two-way ANOVA with Tukey’s multiple comparison test; ns, not significant. See also Figure S5.

In mammals, choline is irreversibly oxidized to betaine in the liver through two sequential reactions catalyzed by choline dehydrogenase and betaine aldehyde dehydrogenase (Ueland, 2011) (Figure 5A). The *C. elegans* genome contains orthologs for both choline dehydrogenase (*chdh-1*) and betaine aldehyde dehydrogenase (*alh-9*) genes (Figure 5A). Like *metr-1*, transcriptional reporters for both *chdh-1* and *alh-9* are expressed in the intestine and hypodermis (Figure 5E and Figure S5B). We found that RNAi knockdown of either *chdh-1* or *alh-9* completely abolished the suppression effect of vitamin B12 on reversals of *unc-2(gf)* mutants (Figure 5F). In fact, RNAi knockdown of *chdh-1* further increased reversals (Figure 5F), similar to the *metr-1* mutation (Figure 4C). *chdh-1* and *alh-9* RNAi did not diminish the effect of vitamin B12 on animal growth of *unc-2(gf)* mutants (Figure S5C), providing further evidence that the effect of vitamin B12 on behavior is independent of its role in animal growth. Together, these results suggest that vitamin B12 modulates behavior through metabolic crosstalk between the Met/SAM cycle and choline oxidation pathway (Figure 5A).

### Betaine can act as a methyl donor in the Met/SAM cycle

In vertebrates, betaine, the product of choline oxidation pathway, and 5-meTHF can act as methyl donors for the synthesis of methionine from homocysteine through the activity of two closely related enzymes: betaine-homocysteine methyltransferase (BHMT) and methionine synthase (METR), respectively (Evans et al., 2002) (Figure 5A). *C. elegans*, like other invertebrates, however, lacks a gene encoding BHMT (Wasmuth et al., 2008). This raised the possibility that the invertebrate methionine synthases can use both betaine and 5-meTHFas a methyl donor. We analyzed the *in vivo* interaction between betaine and Met/SAM cycle. Betaine supplementation, like methionine or SAM supplementation, increased animal growth in a dose-dependent manner (Figure 6A and Figure S4G). However, betaine supplementation failed to increase animal growth of *metr-1* mutants (Figure 6B). In mammals, betaine and 5-meTHF can complement each other to maintain flux through the Met/SAM cycle (da Costa et al., 2005; Ueland, 2011). Consequently, in mice, betaine supplementation can rescue the early postnatal mortality caused by 5-meTHF deficiency of methylenetetrahydrofolate reductase (*Mthfr*) knockout mice (Schwahn et al., 2004). Interestingly, polymorphisms in the MTHFR gene in humans increases susceptibility to migraines (Azimova et al., 2013; Liu et al., 2014). In *C. elegans*, deletion of *mthf-1*/*MTHFR* results in 5-meTHF deficiency and developmental arrest at early larval stages (Maydan et al., 2007). We found that betaine supplementation significantly rescued the larval arrest of *mthf-1* mutants: Only ∼10% of *mthf-1* mutants reached adulthood on *E. coli* OP50 diets, whereas ∼ 40% of *mthf-1* homozygous mutants reached adulthood when betaine was supplemented to *E. coli* (Figure 6C). Vitamin B12 plus betaine further rescued the development of *mthf-1* mutants (Figure 6D). Thus, betaine, the product of the choline oxidation pathway, acts as methyl donor in the Met/SAM cycle.

**Figure 6.**
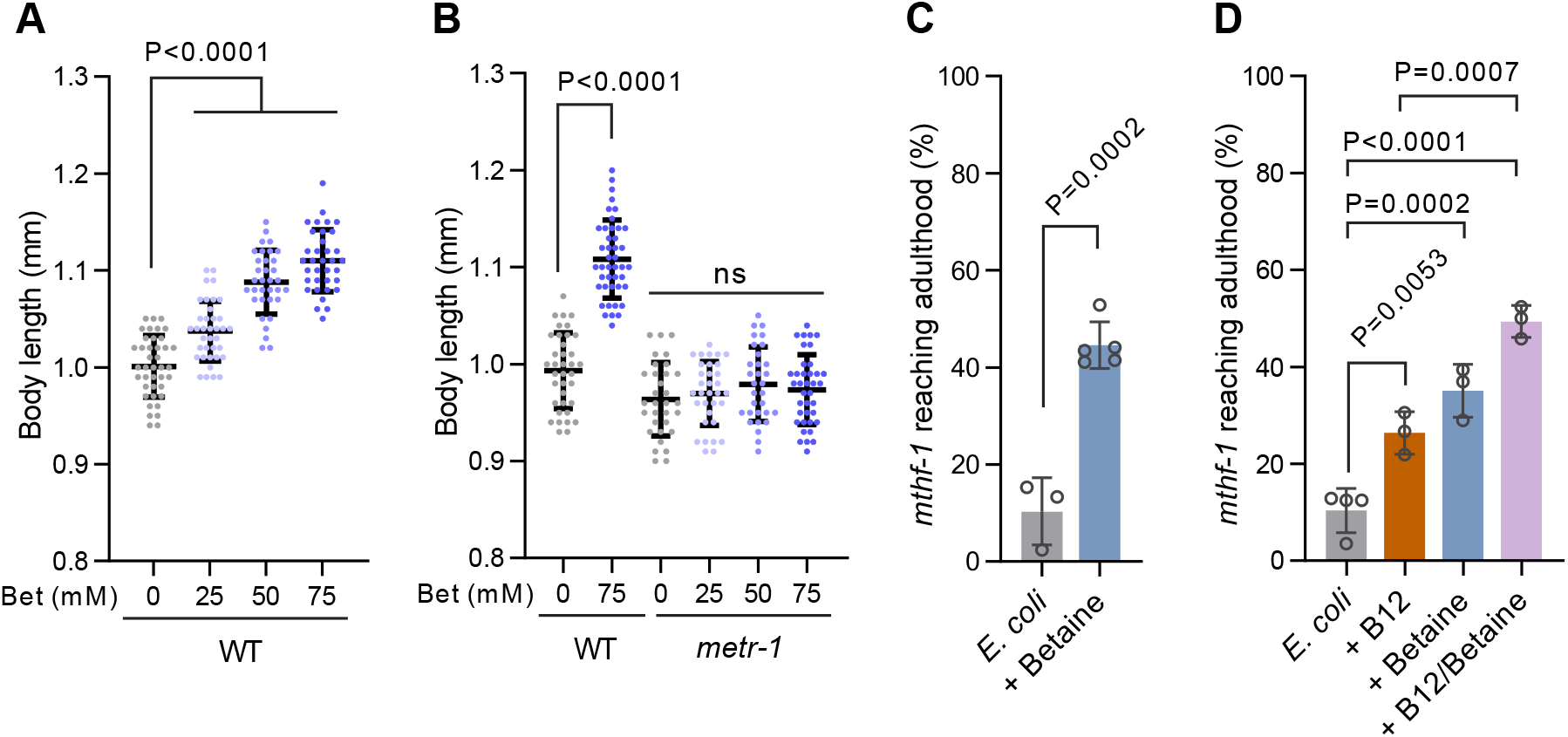
Betaine acts as a methyl donor in the Met/SAM cycle. (A) Growth rate as indicated by body length of wild-type fed *E. coli* with the indicated concentrations of betaine. (B) Growth rate as indicated by body length of wild-type and *metr-1* mutants fed *E. coli* with the indicated concentrations of betaine. (C) Percentage of *mthf-1; bli-2* mutants that reached adulthood on *E. coli* with or without 75 mM betaine. (D)Percentage of *mthf-1; bli-2* mutants that reached adulthood on *E. coli, E. coli* with B12 (64 nM), *E. coli* with betaine (75 mM), or *E. coli* with both B12 and betaine. All data are presented as mean ± s.e.m. from at least three independent experiments. P-values are from two-way ANOVA with Tukey’s multiple comparison test; ns, not significant.

### A neuronal choline transporter is required to mediate the effect of B12 on excitatory transmission

Crosstalk between the choline oxidation pathway and the Met/SAM cycle can impact levels of free choline (Rosas-Rodriguez and Valenzuela-Soto, 2021; Ueland, 2011). Choline supply for acetylcholine biosynthesis in cholinergic neurons occurs in part through the uptake of free choline by high-affinity choline transporters. Choline uptake is modulated to meet the demands of increased ACh synthesis and release (Simon and Kuhar, 1975). Choline transporter (CHT) mouse knockouts have severe defects in cholinergic transmission resulting in early neonatal lethality indicating that choline uptake through CHT is a major source of choline for ACh biosynthesis. *C. elegans* has a single choline transporter, CHO-1 (Matthies et al., 2006). *cho-1* null mutants are viable, display normal locomotion on agar plates but fail to sustain swimming behavior in liquid (Mullen et al., 2007). Loss of *cho-1* function reduced reversals and aldicarb sensitivity of *unc-2(gf)* mutants (Figures 7A and 7B). This result suggests that free choline and choline transport is required to maintain elevated levels of cholinergic transmission in *unc-2(gf)* mutants. Therefore, we next tested whether choline uptake by CHO-1 in cholinergic neurons is required for the effect of vitamin B12 on behavior. Vitamin B12 supplementation did not further suppress the reversals and aldicarb sensitivity of *cho-1; unc-2(gf)* mutants (Figures 7A and 7B) but still stimulated animal growth (Figure S6). These results support a gut-brain communication pathway in which vitamin B12-dependent metabolism in the intestine reduces excitatory cholinergic signaling in the nervous system by decreasing the availability of free choline required for the synthesis of acetylcholine (Figure 7C).

**Figure 7.**
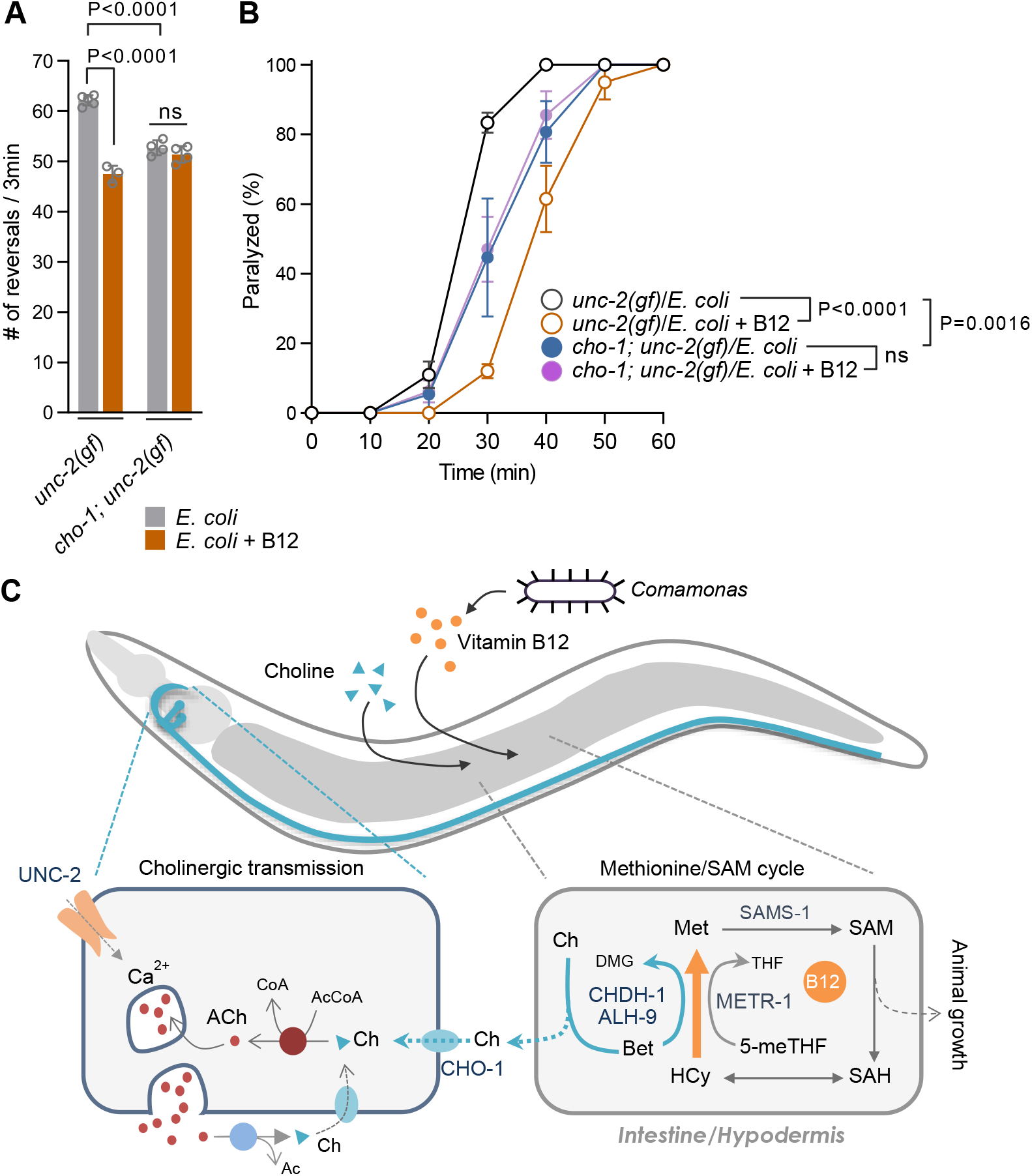
A neuronal choline transporter is required to mediate the effect of B12 on excitatory transmission. (A) Reversal frequency of *unc-2(gf)* and *cho-1; unc-2(gf)* mutants fed *E. coli* with or without 64 nM B12. (B) Quantification of paralysis percentage of *unc-2(gf)* and *cho-1; unc-2(gf)* mutants fed *E. coli* with or without 64 nM B12 on 1 mM aldicarb. (C) Model: Vitamin B12 produced by gut bacteria modulates excitatory neurotransmission of the host *C. elegans*. Metabolic crosstalk between the B12 dependent Met/SAM cycle and the choline-oxidation pathway in the intestine and hypodermis lowers the levels of free choline. The reduced choline availability by B12 limits acetylcholine synthesis in the neurons particularly under conditions of elevated acetylcholine release. All data are presented as mean ± s.e.m. from at least three independent experiments. P-values are from two-way ANOVA with Tukey’s multiple comparison test; ns, not significant. See also Figure S6.

## DISCUSSION

Diet and gut microbiota play a critical role in neuronal health and disease. Elucidating the impact of individual nutrient and microbiota on host nervous system function and its underlying mechanisms is challenging and our understanding is still at an early stage (Cryan et al., 2019; Fischbach, 2018; Walter et al., 2020). The complexity of the mammalian diet, microbiome and nervous system have limited the ability to define causal relationships between the microbiota and behavior. Here, we leveraged the defined bacterial diet of *C. elegans* to identify microbiota that modulate host neural function and behavior. In a survey of bacterial diets, we find that microbiota that produce vitamin B12 and colonize the intestine, lead to persistent changes in *C. elegans* behavior. Our data indicate that vitamin B12 reduces excitatory cholinergic signaling through metabolic cross talk between Met/SAM cycle and choline oxidation pathway.

Vitamin B12 is exclusively synthesized by certain bacteria and archaea (Bender, 2003). *C. elegans* like other animals, must obtain B12 through their diet or symbiosis with bacteria in their intestine (Bito et al., 2013; Kon and Porter, 1954; LeBlanc et al., 2013). In nematodes, B12 regulates development, lifespan, lysosomal activity, gene expression and predatory behaviors (Akduman et al., 2020; McDonagh et al., 2022; Watson et al., 2014; Wei and Ruvkun, 2020). Human intestinal microbiota can also produce vitamin B12 (Albert et al., 1980; Baker, 1981; Hill, 1997; LeBlanc et al., 2013), although the consumption of animal products is likely to be the main source for this nutrient. Vitamin B12 is an essential cofactor for two highly conserved metabolic enzymes methionine synthase and methylmalonyl-CoA mutase (Bito et al., 2013; Kolhouse and Allen, 1977; Watson et al., 2014). Our data show that the B12 dependent reduction of cholinergic signaling depends on methionine synthase, a key enzyme in the Met/SAM cycle. The B12 dependent stimulation of the Met/SAM cycle and the synthesis of methionine and SAM was previously found to accelerate development (Watson et al., 2014). In vertebrates, methionine is synthesized in the Met/SAM cycle from homocysteine by two related enzymes using 5-meTHF (methionine synthase) or betaine (BHMT) as methyl donors. *C. elegans* and other invertebrates lack BHMT and suggesting that they may rely on a single methionine synthase (METR-1). Our finding that betaine stimulates animal growth in a *metr-1* dependent manner is consistent with the hypothesis that METR-1 can use both betaine and 5-meTHF as methyl donors for methionine synthesis. Betaine is synthesized from choline in the choline oxidation pathway. The metabolic connection between Met/SAM cycle and choline oxidation pathway is well established in mammals (da Costa et al., 2005; Ueland, 2011). In *C. elegans*, we find that the choline-oxidation pathway genes *alh-9* and *chdh-1* are co-expressed with *metr-1* in the intestine and the hypodermis, consistent with the metabolic crosstalk between the Met/SAM cycle and the choline oxidation pathway. Crosstalk between the choline oxidation pathway and the Met/SAM cycle impacts levels of free choline, the rate-limiting precursor in the synthesis of acetylcholine. Acetylcholine biosynthesis in cholinergic neurons occurs in part through the uptake of free choline by high-affinity choline transporter CHO-1. We find that the inhibitory effect of vitamin B12 on cholinergic transmission is blocked by mutations in the choline transporter CHO-1. Our findings are consistent with a gut-brain communication pathway in which dynamic crosstalk between vitamin B12-dependent Met/SAM cycle and the choline oxidation pathway decreases the availability of free choline required for the synthesis of acetylcholine (Figure 7C).

The impact of vitamin B12 on cholinergic transmission and behavior is not obvious under normal conditions but becomes apparent under conditions where cholinergic signaling is increased, either genetically (*unc-2(gf), acr-2(gf)*, and *ace-1; ace-2* mutants) or during physical exertion (swimming). Under normal conditions, choline levels in cholinergic neurons can be sustained by re-uptake mechanism at the synapses (Ferguson et al., 2004; Jope, 1979). However, under genetic or behavioral conditions of increased acetylcholine release, reduced choline availability limits acetylcholine synthesis. This is consistent with observations in rodents where the synthesis and release of acetylcholine becomes strongly dependent on choline availability under conditions of increased acetylcholine release (Cohen and Wurtman, 1976; Hartmann et al., 2008; Koppen et al., 1997). Vitamin B12 induced rewiring of choline metabolism may limit acetylcholine synthesis particularly under conditions of elevated acetylcholine release. Our findings provide insight why the relationship between diet, microbiota and the nervous system often only become apparent under conditions where the host is challenged either genetically or environmentally.

Alterations in gut microbiota have been associated with many human diseases (Arzani et al., 2020; Dahlin and Prast-Nielsen, 2019; Golubeva et al., 2017; Lum et al., 2020; Sharon et al., 2019; van Hemert et al., 2014). Human intestinal bacteria also produce vitamin B12 (Albert et al., 1980; Baker, 1981; Hill, 1997; LeBlanc et al., 2013). Vitamin B12 is important for overall health, but particularly for the brain. A wide variety of neurological disorders are also commonly associated with vitamin B12 deficiency (Dror and Allen, 2008; Green et al., 2017; Molloy et al., 2009). Vitamin B12 supplementation has been shown to alleviate the symptoms of several neurological disorders including autism, epilepsy, schizophrenia and migraine (Mitra et al., 2017; Pineles et al., 2010; Shaik et al., 2014). However, the underlying mechanisms of the positive impact of B12 are largely unclear. Notably, these neural disorders have been associated with a shift in excitatory and inhibitory balance toward excitation in the nervous system (Eichler and Meier, 2008; Nelson and Valakh, 2015; Vecchia and Pietrobon, 2012). As acetylcholine is a main excitatory neurotransmitter in the brain, it will be interesting to determine whether vitamin B12 dampens excitatory signaling in the nervous system and thus improves the symptoms of neurological disorders that are associated with excitatory and inhibitory imbalance.

## Supporting information

Supplemental fugures_Kang et al

## ACKNOWLEDGMENTS

We thank the Caenorhabditis Genetics Center (CGC), which is funded by the NIH Office of Research Infrastructure Programs (P40 OD010440), for some worm and bacterial strains. We thank B. Samuel, H. Schulenburg, M. Shapira, and V. Ambros for bacterial strains; V. Budnik, A. Byrne, A. Walker and C. Vance for helpful discussions; and W. Joyce and J. Florman for technical support. This work was supported in part by National Institutes of Health grant R01 NS107475 (MJA), DK068429 (AJMW), 1R35GM131877 (FCS) and a grant from the Riccio Fund for Neuroscience (MJA and AJMW). FCS is a Faculty Scholar of the Howard Hughes Medical Institute.

## AUTHOR CONTRIBUTIONS

WKK, AB, AT, BWF performed the experiments and analyzed the data. WKK, BWF, FCS, AJMW, MJA designed the experiments. WKK and MJA conceived the study and wrote the paper.

## DECLARATION OF INTERESTS

The authors declare no competing interests.

## MATERIALS and METHODS

### KEY RESOURCES TABLE

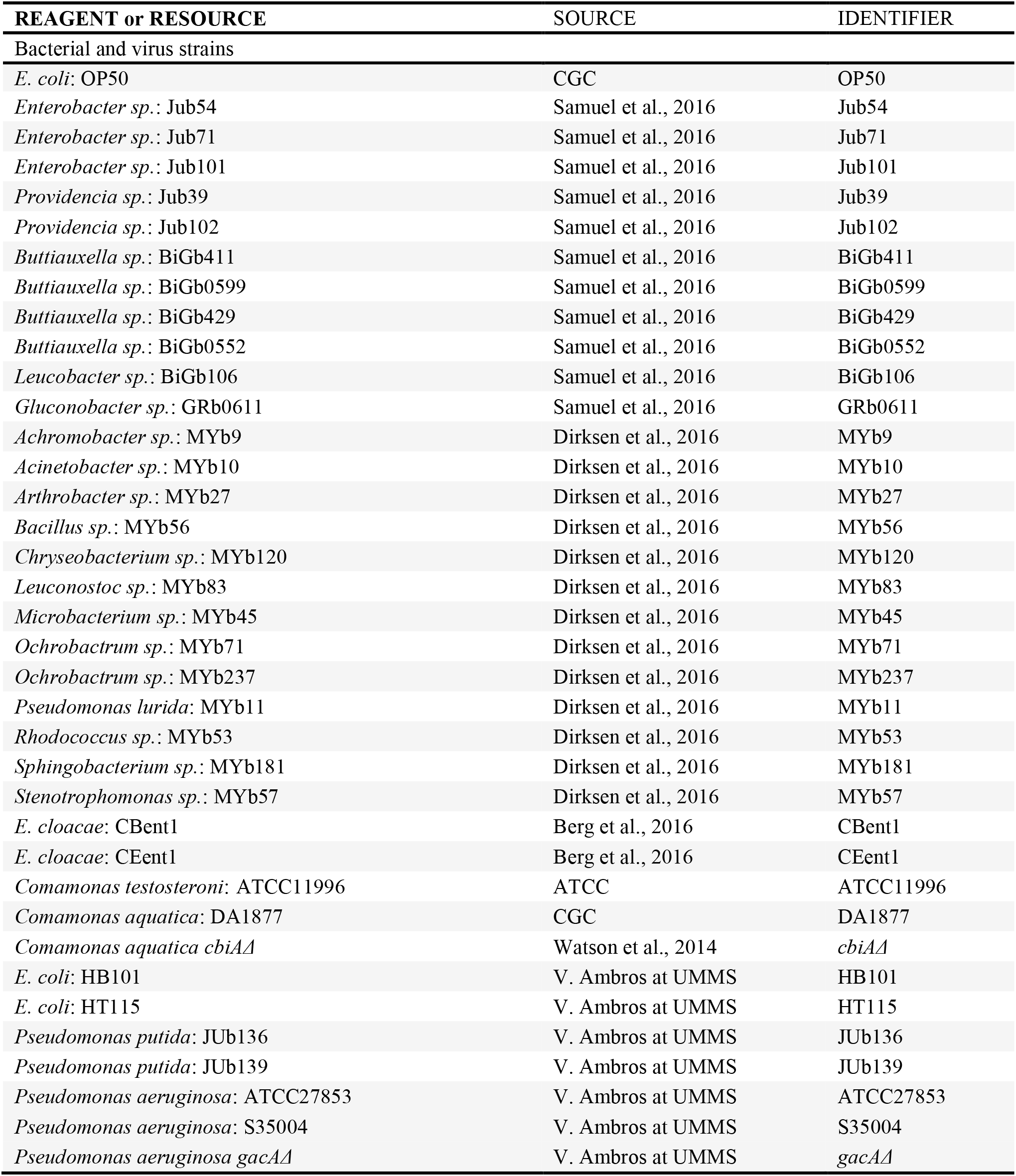
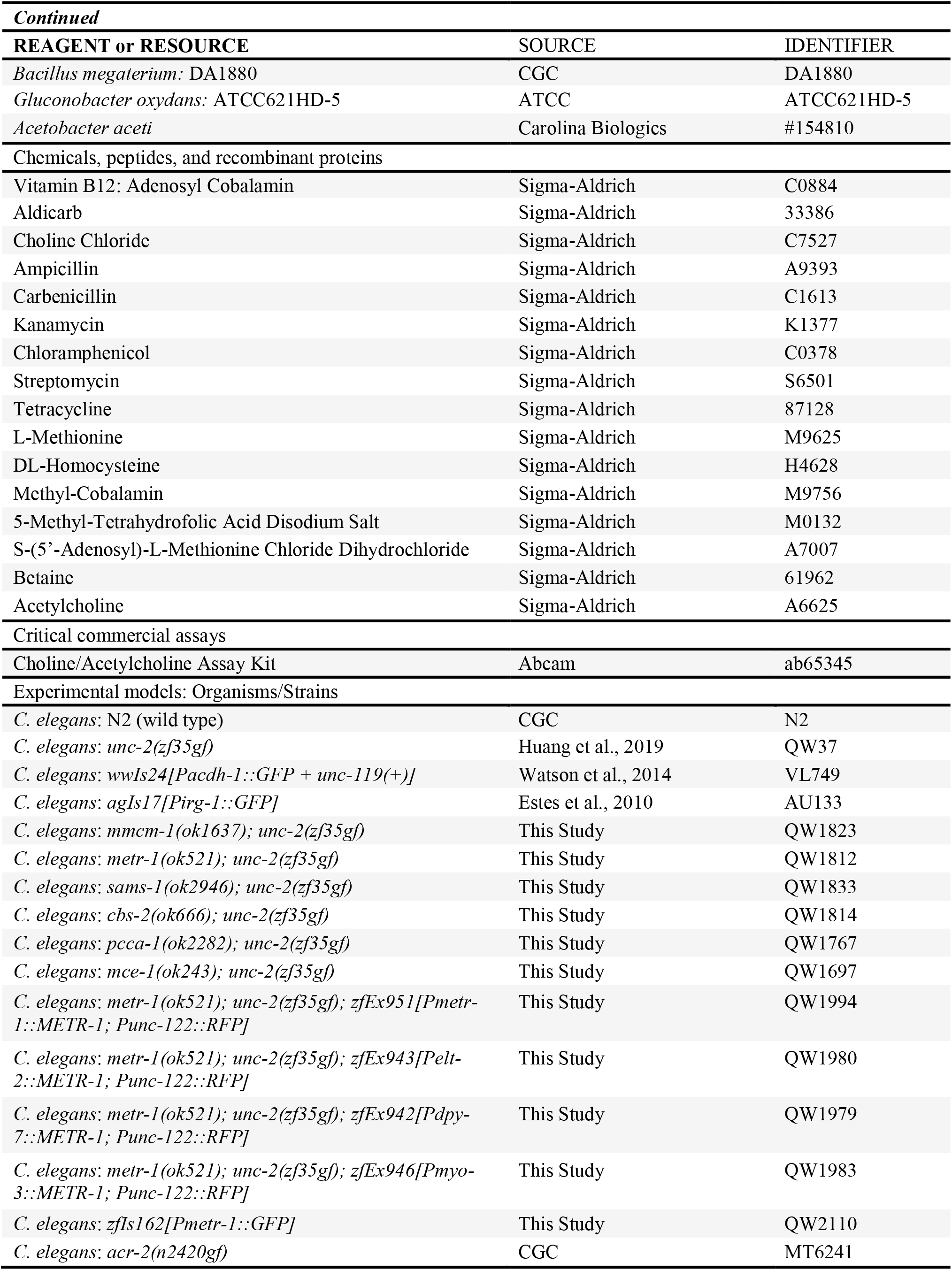
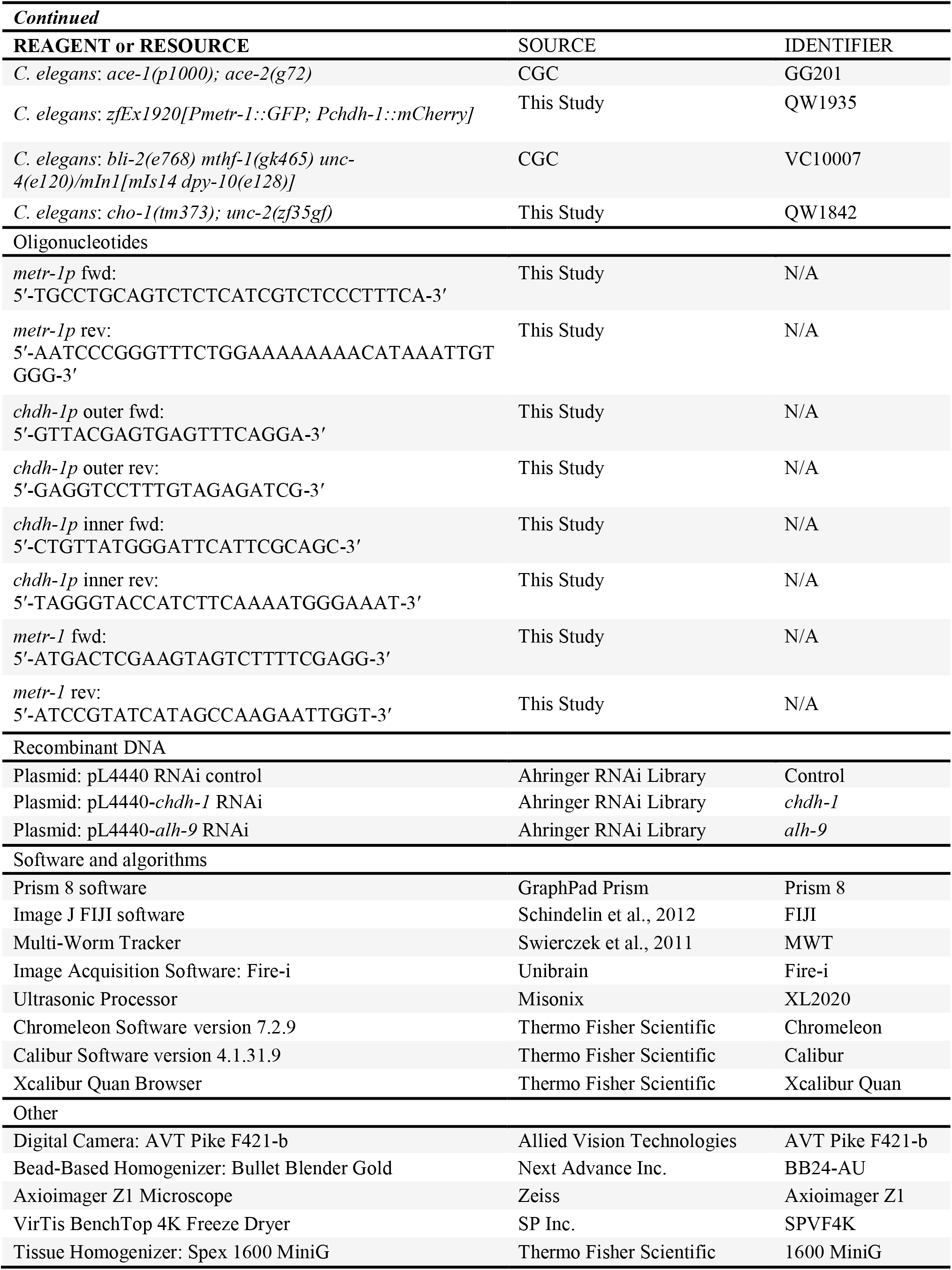

### METHOD DETAILS

#### Nematode strains

All *C. elegans* strains were maintained at 22ºC on nematode growth medium (NGM) plates seeded with *Escherichia coli* OP50 bacteria. The wild-type strain was Bristol N2. MT6241, GG201 and VC10007 strains were obtained from the *Caenorhabditis Genetics Center* (CGC). Transgenic strains were generated by standard microinjection of the desired transgenes and a fluorescent co-injection marker (*Punc-122::RFP*), and at least two independent lines were tested.

#### Bacterial strains

The *E. coli OP50* strain was used as the standard laboratory diet for *C. elegans*. Diverse species of bacterial strains that have been isolated as a natural diet or microbiota of *C. elegans* were kindly provided by *B. Samuel, H. Schulenburg, M. Shapira, M. Walhout* and *V. Ambros*. Some bacterial strains were obtained from the CGC, ATCC, and Carolina Biologics. All bacterial strains were grown overnight at 30ºC in LB media (or mannitol media for *Gluconobacter* and *Acetobacter*). To kill bacteria, an overnight culture of *E. coli* OP50 (or *C. aquatica*) grown in LB was pelleted, washed, concentrated ten-fold in water and incubated at 85°C for 30 min (1h for *C. aquatica*). Absence of live bacteria was confirmed by failure to grow on LB agar plates at 37ºC overnight.

#### Bacterial screen

All behavioral analysis was performed with young adult animals (18-24 h post-L4) at room temperature (22-24ºC). L4 animals grown on *E. coli* OP50 were transferred to plates seeded with the indicated bacterial strain. After 24 h, 20 young adult animals were transferred to an unseeded NGM plate and allowed to crawl away and remove remaining bacteria. The animals were then transferred to a 60-mm NGM plate with a thin lawn of *E. coli* OP50 and allowed to acclimate for 100s. After the acclimation period, the animals were tracked and analyzed using a Multi-Worm Tracker (Swierczek et al., 2011) for 3 min. Videos were recorded at 30 frames per second using a digital camera (AVT Pike F421-b, Allied Vision Technologies) and Fire-i image acquisition software (Unibrain). The number of reversals per animal was calculated as the total number of reversals of the population divided by the average number of animals tracked. Analysis was limited to the animals that had been tracked for a minimum of 20 seconds and had moved a minimum of 2 body lengths.

#### Locomotion assay of *ace-1; ace-2* mutants

L4 animals grown on *E. coli* OP50 were transferred to plates seeded with *E. coli* OP50 plus or minus 64 nM vitamin B12. After 24 h, a single young adult animal was placed on a thin lawn *E. coli* OP50 plate and allowed to acclimate for 100s. Total distance and duration travelled during a forward run was analyzed for 5 min using a Multi-Worm Tracker. Locomotion rate was calculated by distance traveled over time.

#### Convulsion phenotype of *acr-2(gf)* mutants

L4 animals grown on *E. coli* OP50 were transferred to plates seeded with *E. coli* OP50 plus or minus 64 nM vitamin B12. After 24 h, a single young adult animal was placed on a thin lawn *E. coli* OP50 plate and allowed to acclimate for 3 min. Convulsion phenotype of the animal was then manually scored during 1 min as simultaneous contraction of both anterior and posterior body wall muscle.

#### Aldicarb resistance

L4 animals grown on *E. coli* OP50 were transferred to plates seeded with *E. coli* OP50 plus or minus 64 nM vitamin B12. After 24 h, 20 young adult animals were placed in the center of an unseeded NGM agar plate supplemented with 1 mM aldicarb (Sigma-Aldrich). Animals were scored every 10 or 20 min as paralyzed if they did not move when gently touched with a platinum wire on the tail. Animals that crawled off the plate during the analysis were discarded from the analysis.

#### Quiescence phenotype

L4 animals grown on *E. coli* OP50 were transferred to plates seeded with *E. coli* OP50 plus or minus 64 nM vitamin B12 and 30 mM choline. After 24 h, 10 young adult animals were transferred into 24-well plates containing 500 µl M9 buffer in each well. Animals were manually scored every 5 min during 3 h with a dissecting microscope as quiescent if they did not move and maintained a straight rather than sinusoidal posture.

#### Body length measurement

The effect of bacterial diets or metabolites on animal growth was examined by measuring worm body length. L4 animals grown on *E. coli* OP50 were transferred to the plates seeded with the indicated bacterial strains with or without metabolites and grown for 24 h. 20 young adult animals were imaged while freely moving on a thin lawn plate with a digital camera (AVT Pike F421-b) using Fire-I image software (Unibrain). Their body length was quantified using Image J FIJI software.

#### Antibiotic susceptibility test for *C. aquatica*

An overnight culture of *E. coli* OP50 or *C. aquatica* grown in LB was pelleted, washed, and normalized to an OD_600_ of 1.0 with fresh LB. water and incubated at 85°C for 30 min (1h for *C. aquatica*). Normalized cultures were next diluted 1:100 in 1 ml LB media dosed with one of seven antibiotics (ampicillin, carbenicillin, kanamycin, chloramphenicol, streptomycin, and tetracycline) at 1x, 2x, 5x and 10x working concentrations. Cultures were grown at room temperature with gentle rotation for 24 hours, then 10 uL of each culture were plated on LB agar plates. Bacterial colonies were imaged after an overnight incubation at 37ºC.

#### Bacterial colonization assay

To determine bacterial colonization of *C. elegans*, L4 animals grown on *E. coli* OP50 were transferred to plates seeded with *E. coli* OP50, *P. aeruginosa* PA14, or *C. aquatica*. After one day, 10-20 young adult animals were picked and incubated in 1/50 dilution of bleaching solution at RT for 10 min to remove bacteria from the worm surface. The animals were washed 4 times with cold M9, and then lysed using 0.5 mm glass beads with vigorous vortexing in a bead-beater (The Bullet Blender Gold, Next Advance, Inc.) for 3 min at maximum speed. The lysates were diluted in M9 and plated on LB agar plates. Bacterial colonies were counted after an overnight incubation at 37ºC, and the number of bacteria per animal was calculated. Absence of external bacterial contamination was confirmed by plating the last wash on LB agar plates. To evaluate the persistence of colonization, young adult animals grown on *C. aquatica* were transferred to the plates seeded with *E. coli* OP50. After 1, 2, 3, or 4 days, 10-20 animals were picked, washed, lysed, and plated on LB agar plates as described above, and the colonies of *E. coli* OP50 and *C. aquatica* were separately counted according to their distinguishable colony size and morphology. To kill *C. aquatica* bacteria in the worm gut, young adult animals grown on *C. aquatica* were transferred to the plates seeded with *E. coli* OP50 + 200 µg/ml kanamycin. After 4, 8, 12, 24 h, 10-20 animals were picked, washed, lysed, and plated on LB agar plates as described above. The colonies of *C. aquatica* were counted after 24 h to determine if all live bacteria were removed from the worm intestine.

#### Imaging and fluorescence quantification

To test the effects of individual bacterial strains on *acdh-1* expression or immune response of *C. elegans*, L4 animals expressing *Pacdh-1::GFP* (Arda et al., 2010) or *Pirg-1::GFP* (Estes et al., 2010) grown on *E. coli* OP50 were transferred to the plates seeded with a different bacterial strain. After 24 hr, young adult animals were mounted on 2% agarose pads containing 60 mM sodium azide. The GFP images were acquired with an Axioimager Z1 microscope (Zeiss) at 10x or 20x magnification, and the maximum fluorescence intensity was quantified using Image J FIJI software. To determine expression pattern of *metr-1* or *chdh-1*, young adult animals expressing either or both *Pmetr-1::GFP* and *Pchdh-1::mCherry* grown on the plates seeded with *E. coli* OP50 were imaged as described above.

#### Molecular biology

Standard molecular biology techniques were used. For *Pmetr-1::GFP* transcriptional reporter constructs, a 2883 bp promoter fragment upstream of the metr-1 start site was amplified from genomic DNA by PCR and cloned into pPD95.70 GFP plasmid. The plasmid was injected into N2 animals at a concentration of 80 ng/µl and integrated by X-ray irradiation followed by outcrossing three times. For *Pchdh-1::mCherry* transcriptional reporter constructs, a 1200 bp promoter fragment upstream of the chdh-1 start site was amplified from genomic DNA by nested PCR using two pairs of primers and cloned into mCherry vector. The *Pchdh-1::mCherry* plasmid was injected at 80 ng/µl with *Pmetr-1::GFP* (80 ng/µl) into N2 animals. For rescue constructs, the *metr-1* cDNA was amplified from a first-strand cDNA library and cloned behind the endogenous *metr-1* promoter, intestinal *elt-2* promoter, hypodermal *dpy-7* promoter, or muscle-specific *myo-3* promoter. The plasmid was injected at 5 ng/µl with the co-injection marker *Punc-122::RFP* plasmid (80 ng/µl) and pBSK plasmid (75 ng/µl) into *metr-1(ok521); unc-2(zf35gf)* mutant animals.

#### Measurement of free choline and acetylcholine levels

Approximately 25,000 synchronized young adult animals grown in liquid S-complete medium with *E. coli* OP50 plus or minus 64 nM vitamin B12 were washed, resuspended in 10% TCA, and frozen in dry ice/ethanol bath. The samples were sonicated for 90 sec (15 sec on/60 sec off pulse cycles at 20% power) on ice-water bath using ultrasonic processor (Misonix, XL2020) and centrifuged at 16,000 RFC for 30 min at 4ºC. The pellets were resuspended in 1M NaOH and used to determine total protein concentrations using Bradford method (Thermo Scientific, Coomassie Protein Assay Reagent, #1856209). The supernatant was extracted with ether to remove TCA, desiccated, and resuspended in assay buffer. The levels of free choline and acetylcholine were determined using a fluorometric Choline/Acetylcholine assay kit (Abcam, ab65345) according to the manufacturer’s protocol. The values were normalized to total protein concentrations.

#### Sample preparation for HPLC-MS

Worm pellets were lyophilized for 18–24 hr using a VirTis BenchTop 4K Freeze Dryer. Dried pellets were transferred to 1.5 mL microfuge tubes and dry pellet weight recorded. Pellets were disrupted in a Spex 1600 MiniG tissue grinder after addition of three stainless steel grinding balls to each sample. Microfuge tubes were placed in a Cryoblock (Model 1660) cooled in liquid nitrogen, and samples were disrupted at 1100 RPM for six cycles of 30 s. Pellets were transferred to 8 mL glass vials in 6 mL LC-MS grade methanol. Samples were stirred overnight at room temperature following the addition of a stir bar (VWR 58948-375). Glass vials were centrifuged at 2750 RCF for five minutes in an Eppendorf 5702 Centrifuge using rotor F-35-30-17. 5 mL of the resulting supernatant was transferred to a clean 8 mL glass vial and concentrated to dryness in an SC250EXP Speedvac Concentrator coupled to an RVT5105 Refrigerated Vapor Trap (Thermo Scientific). The residue was suspended in 150 µ L methanol, briefly sonicated, transferred to a 1.5 mL microfuge tube and spun at 20,000 RCF in an Eppendorf 5417R Centrifuge for five minutes to sediment precipitate. The resulting supernatant was transferred to an HPLC vial and stored at −20°C or analyzed immediately.

#### HPLC-MS Analysis

Normal-phase chromatography was performed using a Vanquish LC system controlled by Chromeleon Software (version 7.2.9 Thermo Fisher Scientific) and coupled to an Orbitrap Q-Exactive High Field mass spectrometer controlled by Xcalibur software (version 4.1.31.9 Thermo Fisher Scientific). Methanolic extracts prepared as described above (injection volume: 1 µ L) were separated on an Agilent Zorbax RRHD HILIC Plus column (150 mm x 2.1 mm, particle size 1.8 µm) maintained at 40 °C with a flow rate of 0.5 mL/min. Solvent A: 0.1% formic acid in water; solvent B: 0.1% formic acid in acetonitrile. A/B gradient started at 95% B for 2 min after injection and decreased linearly to 50% B at 20 min, followed by a linear decrease to 10% B at 22 min, held at 10% B until 25 min, followed by a linear increase to 90% B at 27 min, and finally held at 90% B for an additional 3 min to re-equilibrate the column. Mass spectrometer parameters: spray voltage (+3.5 kV), capillary temperature 380 °C, probe heater temperature 400 °C; sheath, auxiliary, and sweep gas 60, 20, and 1 AU, respectively. S-Lens RF level: 50, resolution 120,000 at m/z 200, AGC target 3E6, m/z range 70-700. HPLC-MS data were analyzed by integration in Xcalibur Quan Browser (Thermo Fisher Scientific). Acetylcholine and amino acid standards for HPLC-MS were purchased from Sigma Aldrich.

#### RNAi experiments

RNAi bacteria clones obtained from the Ahringer library (Kamath and Ahringer, 2003) were selected by ampicillin (100 mg/ml) and tetracycline (12.5 mg/ml) and verified by sequencing. The RNAi bacteria grown at 37°C overnight in LB with ampicillin (100 mg/ml) were concentrated (4X) and seeded on RNAi NGM plates that contain 6 mM IPTG and 100 mg/ml ampicillin. Synchronized L1 larvae were placed on the RNAi plates and allowed to grow at room temperature. 20 RNAi-treated L4 animals were transferred to new RNAi plates with or without 64 nM vitamin B12. Young adult animals were analyzed for behavioral and growth phenotypes after 24 h.

#### Rescue of developmental arrest caused by *mthf-1* deletion

The balanced heterozygous *mthf-1(gk465)* mutant animals (*bli-2(e768) mthf-1(gk465) unc-4(e120)/mIn1[mIs14 dpy-10(e128)]*), were maintained on *E. coli* OP50 for at least two generations without food deprivation. 4-5 L4 heterozygous *mthf-1(gk465)* mutants were transferred to plates seeded with *E. coli* OP50 plus or minus 64 nM B12 and/or 75 mM betaine. Mutant animals were and allowed to lay eggs for 20-24 h, after which adult animals were removed. The percentage of *Unc mthf-1(gk465)* homozygous mutant animals were followed through development until larval death or reaching the adult stage. WT heterozygous and *Dpy* homozygous animals were removed to avoid food depletion during the assay. *Unc, Bli (bli-2(e768) mthf-1(gk465) unc-4(e120))* mutants that reached to adulthood were scored. Only 10% of *mthf-1(gk465)* homozygous mutants reached adulthood on *E. coli* OP50 diets. In the presence of B12 or betaine, 26% and 35% of *mthf-1(gk465)* homozygous mutants reached adulthood, respectively. 49% of *mthf-1(gk465)* homozygous mutants reached adulthood when supplemented with both B12 and betaine.

#### Quantification and statistical analysis

All data were generated and analyzed using GraphPad Prism 8 software. Results are presented as standard error of the mean (SEM) from at least three independent experiments. Statistical comparisons were performed using ANOVA with Dunnett’s or Tukey’s correction for multiple samples or using an unpaired Student’s t test for two samples. Significance was determined when P-value < 0.05 and the exact P-values were indicated in the relevant graphs. Sample size was chosen based on studies with similar experimental design and was not predetermined by any statistical tests. All experiments were repeated on at least three independent days to ensure reproducibility.

## RESOURCE AVAILABILITY

### Lead contact

Further information and requests for resources and reagents should be directed to and will be fulfilled by the lead contact, Mark J. Alkema.

### Materials availability

All unique reagents including *C. elegans* strains and plasmids etc. generated in this study are available from the Lead Contact without any restriction.

### Data and code availability

All data files are available from the Open Science Framework database: https://osf.io/agpxt/?view_only=0d523c26f9a6497bbb23dd42700a61f6.

## Notes

https://osf.io/agpxt/?view_only=0d523c26f9a6497bbb23dd42700a61f6.

### Competing Interest Statement

The authors have declared no competing interest.

