## Supplemental fugures_Kang et al for "Vitamin B12 produced by gut bacteria modulates excitatory neurotransmission"

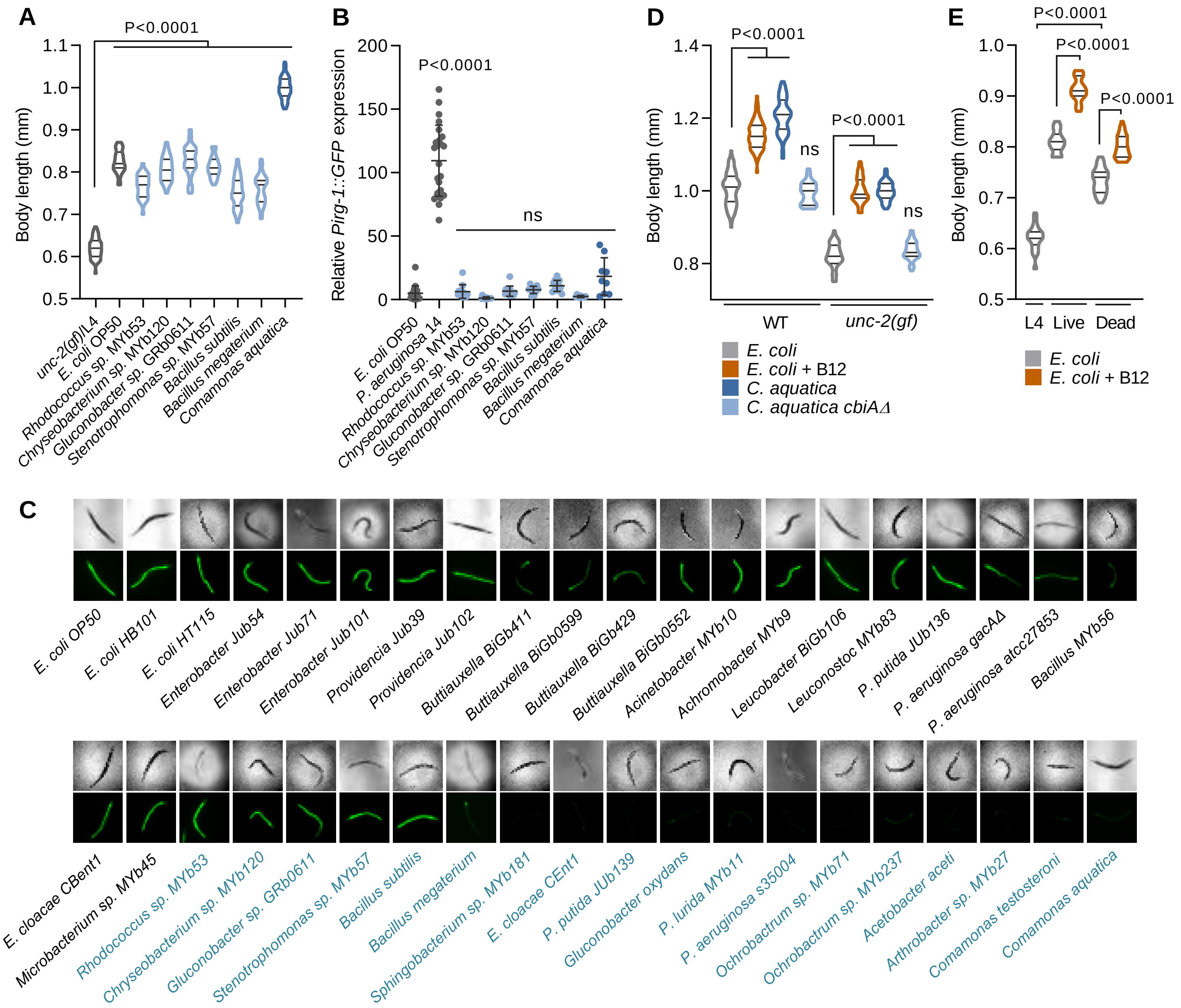

**Figure S1. Effects of different bacterial diets on growth, immune response, and *acdh-1* expression of *C. elegans*, Related to Figure 1**

(A) Growth rate as indicated by body length of *unc-2(gf)* mutants grown on the indicated bacterial strains for 24 h.

(B) Fluorescence values of immune reporter *Pirg-1::GFP* after 24 h exposure to indicated bacterial strains. Pathogenic strain *P. aeruginosa 14* was included as a positive control.

(C) DIC and fluorescence images of *Pacdh-1::GFP* animals fed different bacterial diets for 24 h.

(D) Growth rate as indicated by body length of wild-type and *unc-2(gf)* mutants fed *E. coli*, *E. coli* supplemented with 64 nM B12, *C. aquatica*, or *C. aquatica cbiAΔ* for 24 h.

(E) Growth rate as indicated by body length of wild-type and *unc-2(gf)* mutants fed live or heat-killed *E. coli* with or without 64 nM B12 for 24 h.

All data are presented as mean ± s.e.m. from at least three independent experiments. P-values are from one-way ANOVA with Dunnett's multiple comparison test; ns, not significant.

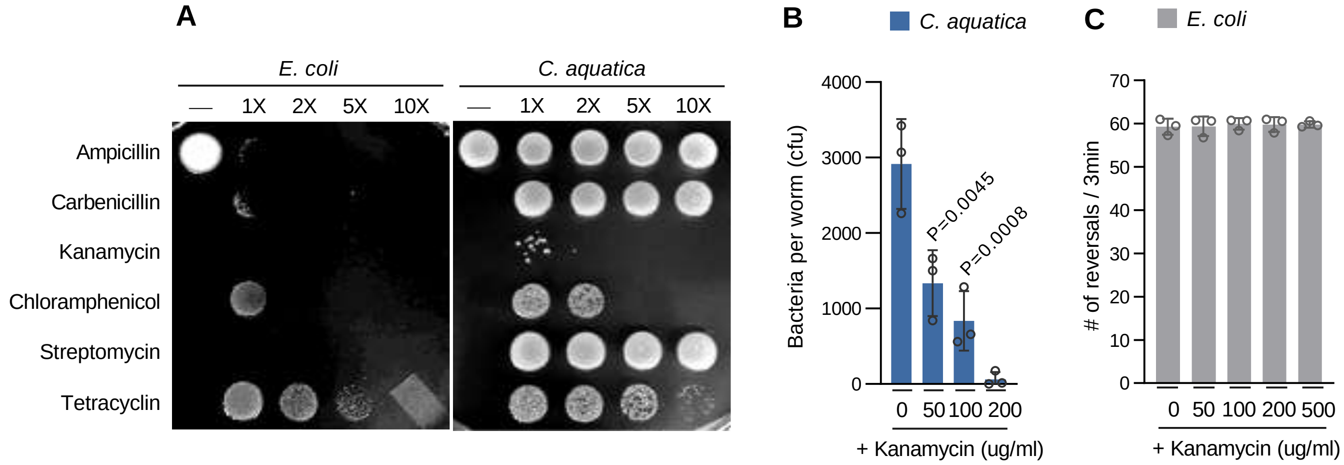


**Figure S2. Antibiotic susceptibility for *C. aquatica* gut bacteria elimination, Related to Figure 2**

(A) Antibiotics susceptibility of *E. coli* and *C. aquatica* treated with the indicated antibiotics for 24 h.

(B) Reversal frequency of *unc-2(gf)* mutants grown on *E. coli* supplemented with indicated concentrations of kanamycin for 24 h.

(C) Bacterial CFU per animal from *unc-2(gf)* mutants grown on *C. aquatica* treated with the indicated concentrations of kanamycin for 24 h.

All data are presented as mean ± s.e.m. from at least three independent experiments. P-values are from one-way ANOVA with Tukey's multiple comparison test; ns, not significant.

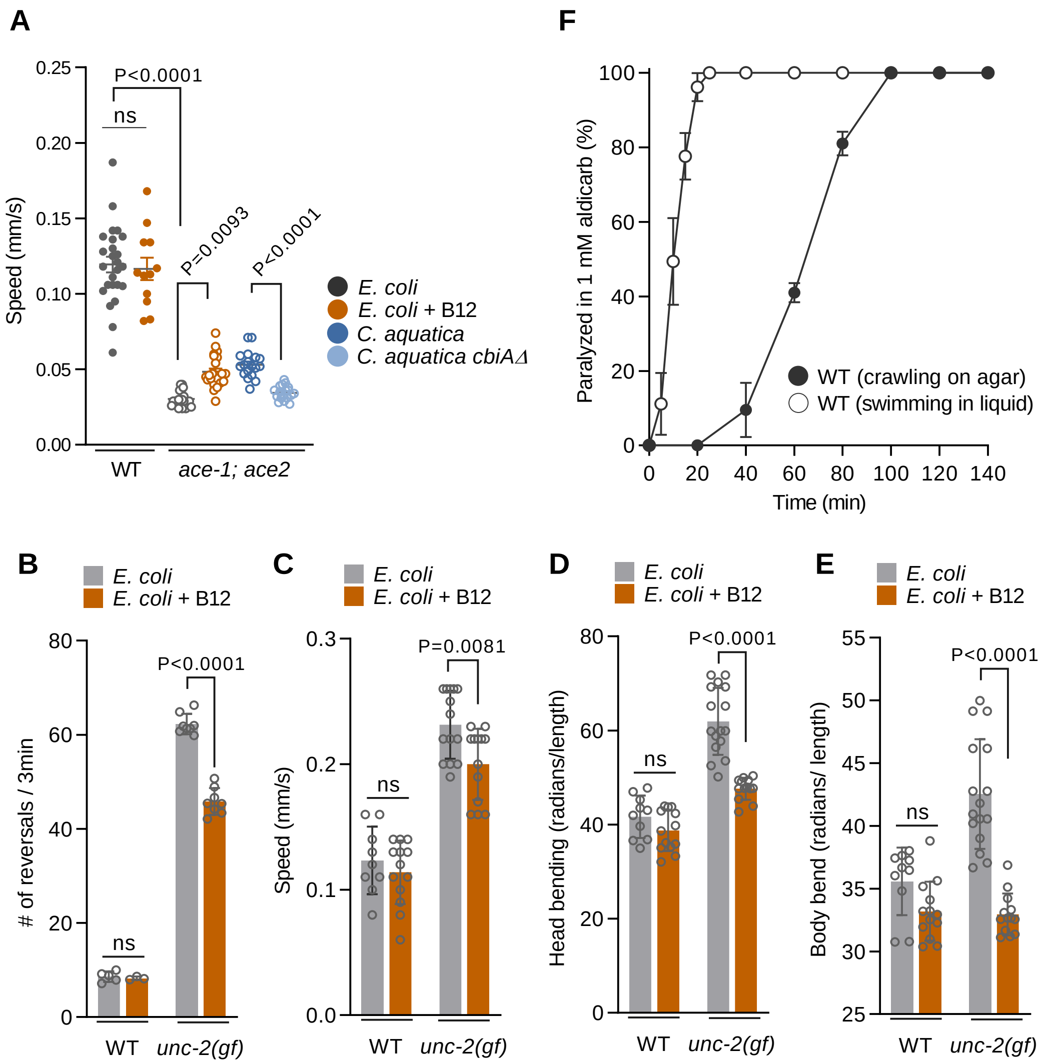


**Figure S3. Vitamin B12 reduces cholinergic signaling especially under conditions of increased acetylcholine release, Related to Figure 3**

(A) Quantification of locomotion speed of wild-type and *ace-1; ace-2* mutants fed *E. coli*, *E. coli* supplemented with 64 nM B12, *C. aquatica*, or *C. aquatica cbiAΔ* for 24 h.

(B-E) Quantification of reversal frequency (B), locomotion speed (C), head bending (D), and body bending (E) of wild-type and *unc-2(gf)* mutants fed *E. coli* with or without 64 nM B12 for 24 h.

(F) Quantification of paralysis percentage of wild-type animals fed *E. coli* on 1 mM aldicarb-containing NGM agar plates or M9 liquid buffer.

All data are presented as mean ± s.e.m. from at least three independent experiments. P-values are from two-way ANOVA with Tukey’s multiple comparison test; ns, not significant.

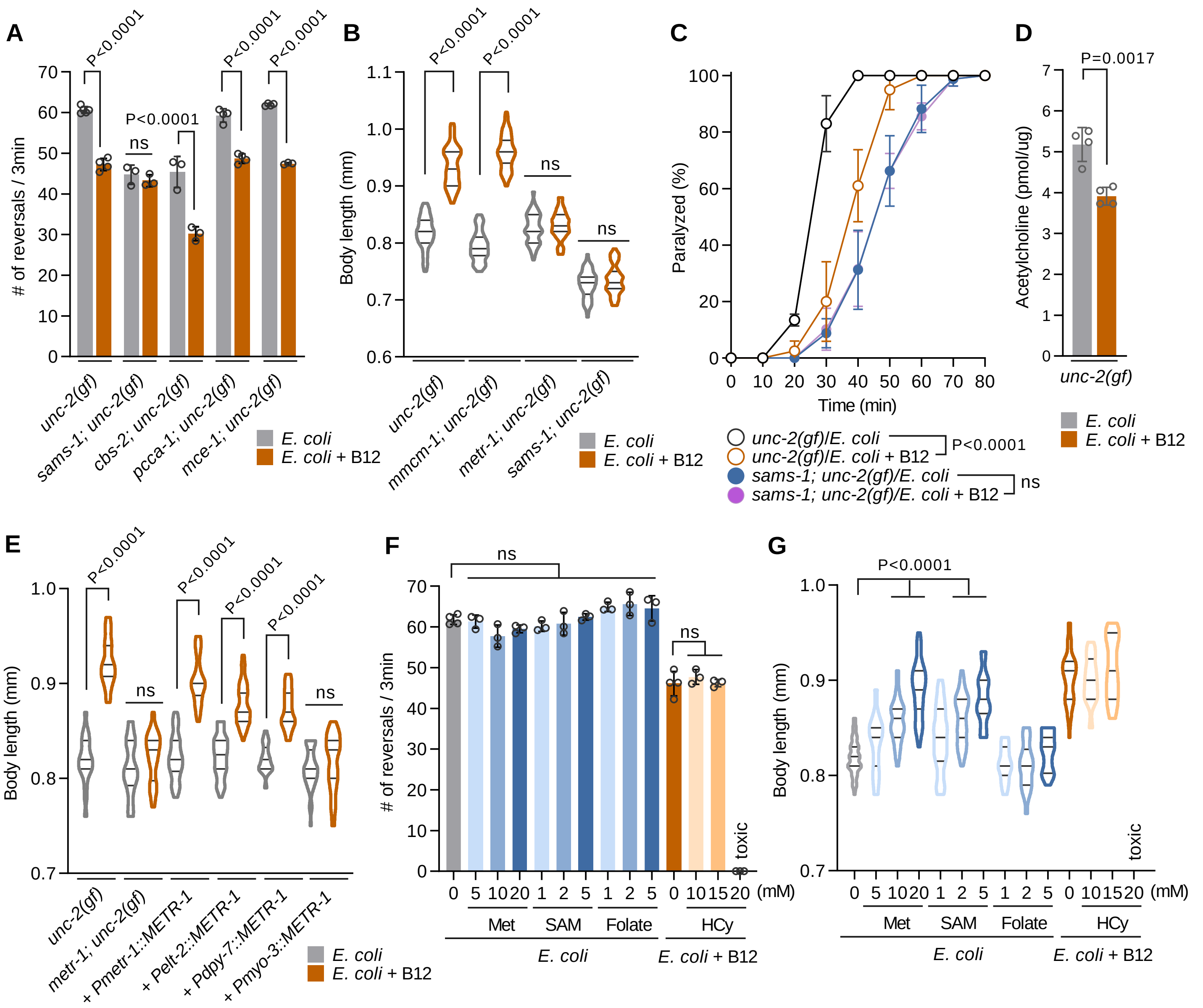


**Figure S4. Vitamin B12 regulates *C. elegans* behavior and growth through Met/SAM cycle, Related to Figure 4**

(A) Reversal frequency of *unc-2(gf), sams-1; unc-2(gf), cbs-2; unc-2(gf), pcca-1; unc-2(gf),* or *mce-1; unc-2(gf)* mutants fed *E. coli* with or without 64 nM B12 for 24 h.

(B) Growth rate as indicated by body length of *unc-2(gf), mmcm-1; unc-2(gf), metr-1; unc-2(gf),* or *sams-1; unc-2(gf)* mutants fed *E. coli* with or without 64 nM B12 for 24 h.

(C) Quantification of paralysis percentage of *unc-2(gf)* and *sams-1; unc-2(gf)* mutants fed *E. coli* with or without 64 nM B12 on 1 mM aldicarb.

(D) Quantification of acetylcholine in *unc-2(gf)* mutants fed *E. coli* with or without 64 nM B12 for 24 h.

(E) Growth rate as indicated by body length of *metr-1; unc-2(gf)* mutants expressing *metr-1* cDNA driven by *Pmetr-1* (endogenous), *Pelt-2* (intestinal), *Pdpy-7* (hypodermal), or *Pmyo-3* (muscle) promoter fed *E. coli* with or without 64 nM B12 for 24 h.

(F) Reversal frequency of *unc-2(gf)* mutants fed *E. coli* supplemented with the indicated metabolites for 24 h.

(G) Growth rate as indicated by body length of *unc-2(gf)* mutants fed *E. coli* supplemented with the indicated metabolites for 24 h.

All data are presented as mean ± s.e.m. from at least three independent experiments. P-values are from one-way ANOVA with Tukey's multiple comparison test; ns, not significant.

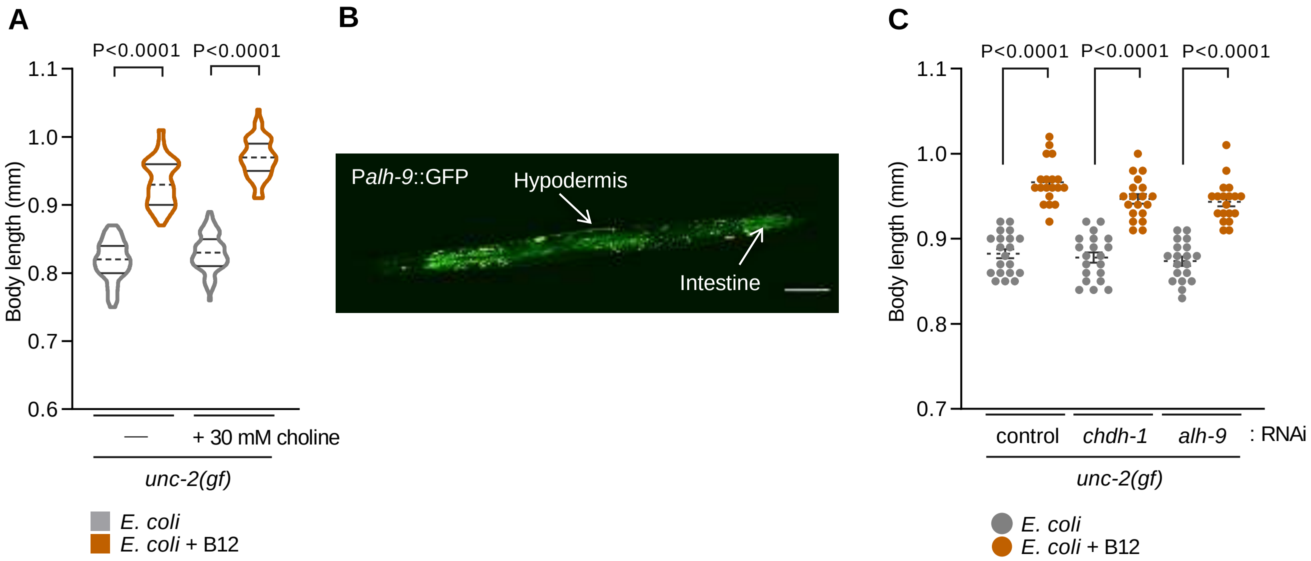


**Figure S5. Choline metabolism is linked to the Met/SAM cycle, Related to Figure 5**

(A) Growth rate as indicated by body length of *unc-2(gf)* mutants fed *E. coli* supplemented with or without 64 nM B12 with 30 mM choline for 24 h.

(B) Expression pattern of *Palh-9::GFP* in the intestine and hypodermis. Scale bar is 100 µm.

(C) Growth rate as indicated by body length of *unc-2(gf)* mutants subjected to RNAi knockdown of *chdh-1* or *alh-9* fed *E. coli* with or without 64 nM B12 for 24 h.

All data are presented as mean ± s.e.m. from at least three independent experiments. P-values are from two-way ANOVA with Tukey's multiple comparison test; ns, not significant.

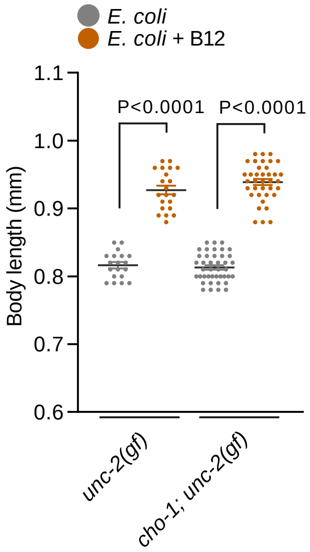


**Figure S6. A neuronal choline transporter is required to mediate the effect of B12 on excitatory transmission, Related to Figure 7**

Growth rate as indicated by body length of *unc-2(gf)* and *cho-1; unc-2(gf)* mutants fed *E. coli* with or without 64 nM B12 for 24 h. All data are presented as mean ± s.e.m. from at least three independent experiments. P-values are from two-way ANOVA with Tukey's multiple comparison test; ns, not significant.
